# Enzyme-responsive Hemostatic Elastin-like Polypeptides for Fibrin Stabilization and Coagulation Restoration in Thrombocytopenia

**DOI:** 10.1101/2025.07.11.664375

**Authors:** Yang Sun, Sunghyun Kang, Malvina Heiniger, Ivan Urosev, Rosario Vanella, Adrian Bertschi, Michael A. Nash

## Abstract

Hemorrhage remains a leading cause of mortality in trauma and surgery, and treatment options are limited for thrombocytopenic patients with impaired platelet function. Current plasma-derived hemostatic products face challenges including limited supply, storage requirements, and infectious risk. Here we report a recombinant protein-based hemostat designed to enhance clot mechanics through enzyme responsiveness and self-assembly, which integrates biophysical design principles with clot-targeted drug delivery. We rationally designed a library of enzyme-responsive glutamine (Q)-containing block elastin-like polypeptides (Q-block-ELPs) that reinforce fibrin clots through phase separation and covalent cross-linking. Q-block-ELPs incorporate glutamine residues within a peptide motif recognized by coagulation factor XIIIa, enabling site-specific grafting into fibrin networks during clot formation. By tuning polymer length, Q-block valency, and lower critical solution temperature (LCST) behavior, we engineered Q-block-ELPs to phase separate at body temperature and integrate into the fibrin architecture. *In vitro,* Q-block-ELPs increase fibrin network density and stiffness. In a thrombocytopenic mouse model, systemic administration reduced blood loss and accelerated clot formation. This strategy delivers a programmable, pathogen-free platform for systemic bleeding control, bridging biophysical protein design with translational hemostatic therapy, addressing an urgent need for platelet-deficient bleeding disorders.

## Introduction

Hemorrhage is a major cause of preventable death, accounting for roughly 40% of trauma-related fatalities and more than a quarter of maternal deaths worldwide.^1^ Although a range of topical hemostatic agents are available for external bleeding, treatment options for internal or non-compressible hemorrhage remain limited.^2^ Currently available systemic hemostatic therapies typically rely on human plasma-derived products such as whole blood, fresh frozen plasma, platelets, and coagulation factor concentrates.^3^ While effective in many cases, these therapies are limited by a reliance on human donors, high costs for collection and processing, short shelf life, stringent storage requirements, and ongoing concerns over viral transmission and immunological complications.^4^

Recently there has been significant interest in the development of nanomedicine-based intravenous hemostatic agents.^5, 6^ However, translating these approaches into the clinic remains challenging due to multifaceted performance requirements among a diverse and complex set of bleeding conditions. Intravenous hemostats should promote clot formation without increasing the risk of unwanted thrombosis. Additionally, issues related to biocompatibility, biodegradability, immunogenicity, acute toxicity, and clearance pose major hurdles for clinical use. A variety of synthetic strategies have been explored to address these challenges, including small-molecule antifibrinolytics such as tranexamic acid,^7^ micro- and nanoparticle-based platforms,^8, 9, 10^ recombinant coagulation factors,^11^ and synthetic polymers^12, 13^.

Previous work has described platelet-like particles (PLPs) composed of soft poly(N-isopropylacrylamide-co-acrylic acid) hydrogels functionalized with fibrin-binding ligands for intravenous hemostasis^9^. Because of their size (∼1 µm), PLPs are rapidly cleared from circulation by macrophages in the liver and spleen through the reticuloendothelial system (RES).^14^ Other nanoparticle-based intravenous hemostatic agents have been developed, including liposomes bearing fibrinogen-derived peptides, ligands targeting von Willebrand factor or platelet glycoprotein IIb/IIIa, and silica nanoparticles conjugated with polyphosphates, all of which have been shown to promote fibrin network formation and enhance clotting.^10, 15^ Despite these advances, nanoparticle-based platforms continue to face challenges related to RES clearance, infusion-related immune activation, and the complexities associated with nano-manufacturing and regulatory approval. Soluble synthetic polymers have also been explored, for example, poly(STAT), a poly(2-hydroxyethyl methacrylate) linear polymer functionalized with fibrin-binding peptides was shown to reinforce fibrin networks and improve survival in rat models of femoral artery hemorrhage.^5, 12, 16^

To date, these materials have primarily been evaluated in hemorrhage models using animals with standard coagulation profiles. However, such materials hold particular relevance for thrombocytopenia, a condition where reduced platelet counts markedly increase bleeding risk following injury^17^. Thrombocytopenia can occur in a variety of clinical settings, including malignancy and immune-mediated disorders. Current therapies often rely on thrombopoietin receptor agonists (TRAs) to stimulate platelet production^18^. Several forms of thrombocytopenia including severe aplastic anemia and subtypes of immune thrombocytopenia are unresponsive to TRA therapy^19^. In addition, TRAs may be associated with an increased risk of thromboembolic events.^20^ Plasma-derived products remain in use for acute hemorrhage control in thrombocytopenic patients. Together, these challenges underscore the need for novel therapeutic strategies that provide safer and more effective control of bleeding in thrombocytopenia and trauma.

Elastin-like polypeptides (ELPs) are a class of synthetic protein polymers inspired by the hydrophobic domain of the human extracellular matrix protein tropoelastin.^21^ They consist of repeating (VPGXG)_n_ pentapeptide motifs, where the guest residue (X) can be substituted with any amino acid excluding proline.^22^ ELPs have been widely explored for applications in drug delivery, tissue engineering, and biosensing^23^ in part because their amino acid sequences and physicochemical properties such as their lower critical solution temperature (LCST) can be precisely controlled by genetic engineering.^24^ Above the LCST, ELPs undergo an entropically driven phase transition, separating into protein-rich coacervates.^25^ The LCST can be adjusted across a broad temperature range by modifying variables such as the guest residue composition and the number of pentapeptide repeats.^26^

In this work, we developed enzyme-responsive ELPs that leverage an enzymatic activity that is spatially and temporally triggered during natural coagulation. Once the coagulation cascade is initiated upon bleeding, a series of proteolytic activation steps leads to the cleavage of factor XIII (FXIII) by thrombin, generating its active transglutaminase form, FXIIIa. FXIIIa catalyzes a Ca^2+^-dependent transamidation reaction between the γ-carboxamide groups of glutamine (Q) residues and the ε-amino groups of lysine (K) residues within fibrin, forming stable covalent isopeptide bonds that cross-link fibrin and strengthen the clot.^27^ To take advantage of this physiological mechanism, we engineered enzyme-responsive ELPs with an LCST slightly below body temperature, leading to phase separation upon administration *in vivo* and formation of particle-like coacervates. To exploit the spatio-temporal control of FXIIIa-mediated transamidation, we introduced peptide sequences containing glutamine (Q) residues embedded within a contextual peptide sequence (DQMMLPWPAVAL) that we call Q-block that is efficiently recognized by FXIIIa.^24, 28^ Upon encountering FXIIIa, Q-blocks are covalently ligated to K-residues in a proximity-dependent manner. Due to the multi-valency of Q-blocks within out ELPs constructs, this mechanism provides a synthetic means for cross-linking fibrin chains within nascent blood clots.

We generated a library Q-block-ELPs with varying Q-block copy numbers at the N-terminus and C-terminus of ELPs of different lengths. This approach allowed us to study sequence-performance relationships for Q-block-ELP coacervates as they covalently integrated into the fibrin network, reinforced the clot structure, and leveraged the natural enzymatic activity of the coagulation cascade. Our work explores the biophysical characteristics of Q-block-ELPs and assesses their ability to regulate coagulation dynamics and blood loss in thrombocytopenic mouse models of hemorrhage.

## Results and Discussion

### Design of Q-block-ELP sequences

We designed a series of ELP constructs using an ABA block architecture (**Figure 1a**). Each block contained 10 or 11 repeats of the VPGXG pentapeptide, where the guest residue (X) consisted of varying proportions of alanine, valine, glutamic acid, and tyrosine. For example, in the ELP1 construct denoted as (Q-block)_4_-(A_2_V_8_E_1_)_2_-(Q-block)_4_ the central block consisted of two repeats of (VPGXG)_11_, in which alanine, valine, and glutamic acid were incorporated at a ratio of 2:8:1, respectively. This central ELP domain was flanked at both the N- and C-termini by four Q-block domains (Q-block), containing four tandem copies of a glutamine-bearing peptide sequence (D<u>Q</u>MMLPWPAVAL)^28^ that is efficiently recognized and cross-linked to K-residues by FXIIIa. Through this mechanism, the Q-block-bearing ELPs can be covalently incorporated into fibrin networks via the coordinated actions of thrombin, calcium, and FXIIIa. The ELP sequences were engineered with LCST values slightly below physiological temperature, allowing them to undergo phase separation upon intravenous administration and form ELP-rich coacervates within the clot (**Figure 1c**). This phase-separated state promotes an enhanced local concentration and may be important for the observed mechanical stability of the resulting clots.

**Figure 1.**
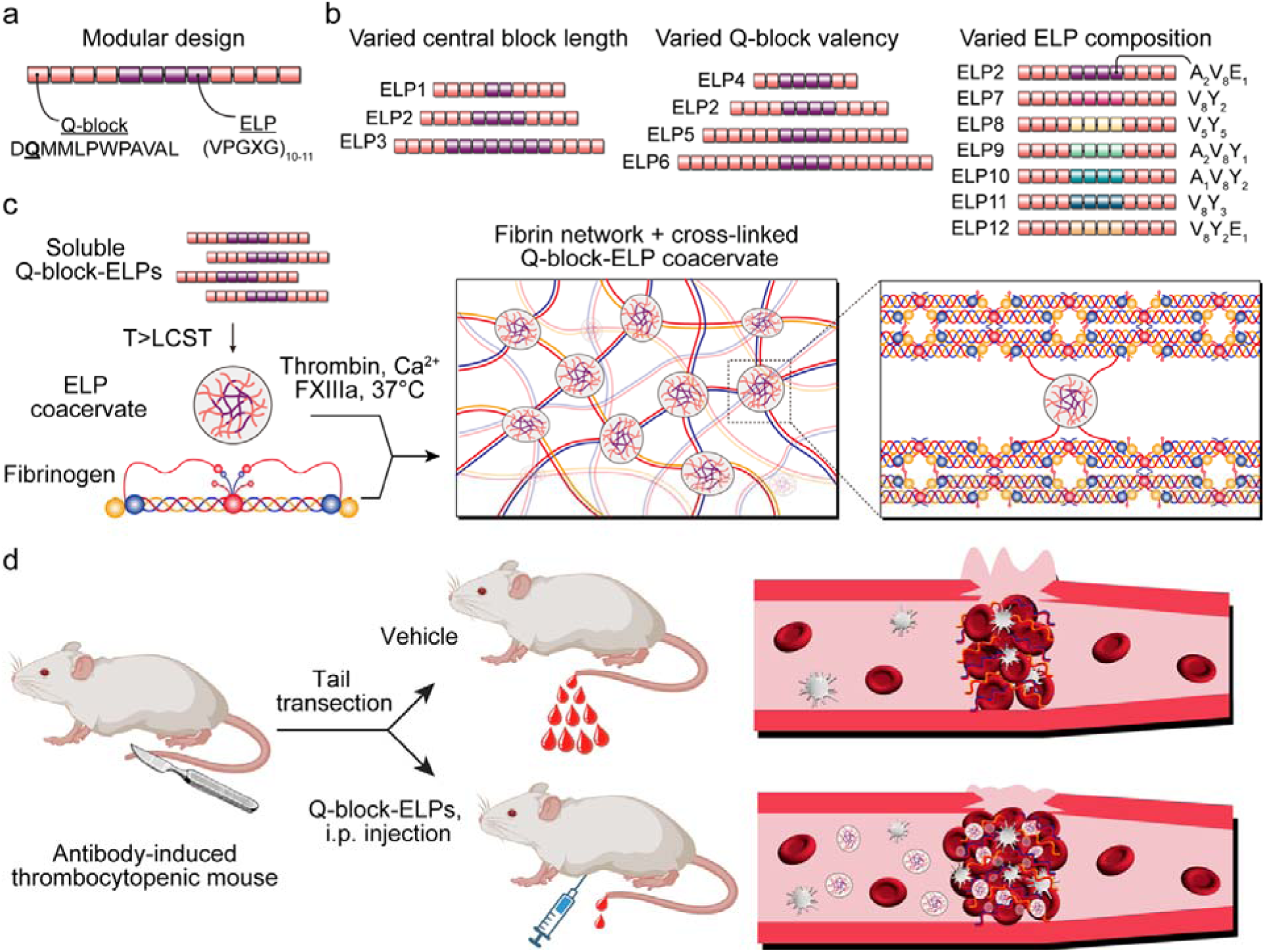
Molecular design and mechanism of Q-block-ELPs for clot reinforcement in thrombocytopenia. (a) Schematic of Q-block-ELP modular architecture. Recombinant elastin-like polypeptides (ELPs) were engineered with glutamine-containing Q-blocks at both termini, flanking a central (VPGXG) repeat domain. The Q-blocks are substrates for the transglutaminase activity of factor XIIIa, enabling site-specific covalent cross-linking into fibrin at locations undergoing active clotting. (b) A library of 12 Q-block-ELPs was constructed by varying central block ELP length (ELPs 1–3), Q-block valency (ELPs 2, and 4-6), and guest residue composition (ELPs 7-12) to systematically explore the influence of sequence features on phase behavior and coagulation. (c) Upon administration at physiological temperature, Q-block-ELPs undergo phase separation and form coacervate droplets that are enzymatically cross-linked into the fibrin network by thrombin-activated FXIIIa, enhancing clot architecture and mechanical resilience. (d) *In vivo* testing in a tail transection bleeding model in thrombocytopenic mice was used to assess efficacy.

To systematically study how the physicochemical properties of ELPs influence fibrin network mechanics, we designed and synthesized a library of 12 ELP constructs that varied in three key parameters: length (central block size), valency (number of Q-block repeats), and hydrophobicity (central block composition) (**Figure 1b**). This design allowed us to examine how adjustments in molecular architecture affect the phase behavior of the ELPs as well as their ability to mechanically reinforce fibrin hydrogels and blood clots. In parallel, we generated a negative control construct (conELP) that shares the same backbone sequence as ELP2, but in which the critical Q-residues were mutated to glycine, thereby preventing FXIIIa-mediated cross-linking.

### Production, Purification, and Phase Separation of Q-block-ELPs

We cloned genes encoding the 12 Q-block-ELP designs (**Table 1**), and expressed and purified the proteins from *E. coli* cultures using inverse transition cycling (ITC) (**Figure 2a**). We measured the LCST of each construct using a spectrophotometric turbidity assay. At a concentration of 30 μM, most ELP constructs exhibited phase transition temperatures below 37°C, indicating that the majority of constructs are aggregated at physiological temperature. As shown in **Figure 2b**, LCST values modestly decreased with increasing length of the central block. Increasing the number of Q-blocks also led to a pronounced reduction in LCST, likely due to greater molecular weight and hydrophobic character of the Q-block, which contains ∼83% hydrophobic amino acids. Substitution of different guest residues at the X-position within the VPGXG motif also influenced LCST values, with more hydrophobic residues resulting in greater reductions in LCST. These findings are consistent with prior reports demonstrating how both chain length and hydrophobicity govern the LCST behavior of ELPs.^21^ Given that hypothermia frequently develops during severe hemorrhage, with core temperatures often dropping to 32-35°C,^29^ tuning the LCST of Q-block-ELPs to remain below these levels may have relevance for maintaining phase separation under clinically relevant conditions of active bleeding.

**Table 1.** Designs of Q-block-ELP constructs. Summary of the 12 ELP constructs generated for this study. Constructs were systematically engineered to vary central block length, Q-block valency, or central block composition. The Q-blocks contain glutamine-rich sequences recognized by FXIIIa, enabling covalent cross-linking into fibrin networks. Sequence notation reflects the guest residue composition of the VPGXG pentapeptide repeats within each construct.

| # | ELP sequence | Design Variable |
| --- | --- | --- |
| ELP1 | (Q-block)<sub>4</sub>-(A<sub>2</sub>V<sub>8</sub>E<sub>1</sub>)<sub>2</sub>-(Q-block)<sub>4</sub> | Central block length |
| ELP2 | (Q-block)<sub>4</sub>-(A<sub>2</sub>V<sub>8</sub>E<sub>1</sub>)<sub>4</sub>-(Q-block)<sub>4</sub> | Central block length |
| ELP3 | (Q-block)<sub>4</sub>-(A<sub>2</sub>V<sub>8</sub>E<sub>1</sub>)<sub>8</sub>-(Q-block)<sub>4</sub> | Central block length |
| ELP4 | (Q-block)<sub>2</sub>-(A<sub>2</sub>V<sub>8</sub>E<sub>1</sub>)<sub>4</sub>-(Q-block)<sub>2</sub> | Q-block valency |
| ELP5 | (Q-block)<sub>6</sub>-(A<sub>2</sub>V<sub>8</sub>E<sub>1</sub>)<sub>4</sub>-(Q-block)<sub>6</sub> | Q-block valency |
| ELP6 | (Q-block)<sub>8</sub>-(A<sub>2</sub>V<sub>8</sub>E<sub>1</sub>)<sub>4</sub>-(Q-block)<sub>8</sub> | Q-block valency |
| ELP7 | (Q-block)<sub>4</sub>-(V<sub>8</sub>Y<sub>2</sub>)<sub>4</sub>-(Q-block)<sub>4</sub> | Central block composition |
| ELP8 | (Q-block)<sub>4</sub>-(V<sub>5</sub>Y<sub>5</sub>)<sub>4</sub>-(Q-block)<sub>4</sub> | Central block composition |
| ELP9 | (Q-block)<sub>4</sub>-(A<sub>2</sub>V<sub>8</sub>Y<sub>1</sub>)<sub>4</sub>-(Q-block)<sub>4</sub> | Central block composition |
| ELP10 | (Q-block)<sub>4</sub>-(A<sub>1</sub>V<sub>8</sub>Y<sub>2</sub>)<sub>4</sub>-(Q-block)<sub>4</sub> | Central block composition |
| ELP11 | (Q-block)<sub>4</sub>-(V<sub>8</sub>Y<sub>3</sub>)<sub>4</sub>-(Q-block)<sub>4</sub> | Central block composition |
| ELP12 | (Q-block)<sub>4</sub>-(V<sub>8</sub>Y<sub>2</sub>E<sub>1</sub>)<sub>4</sub>-(Q-block)<sub>4</sub> | Central block composition |

**Figure 2.**
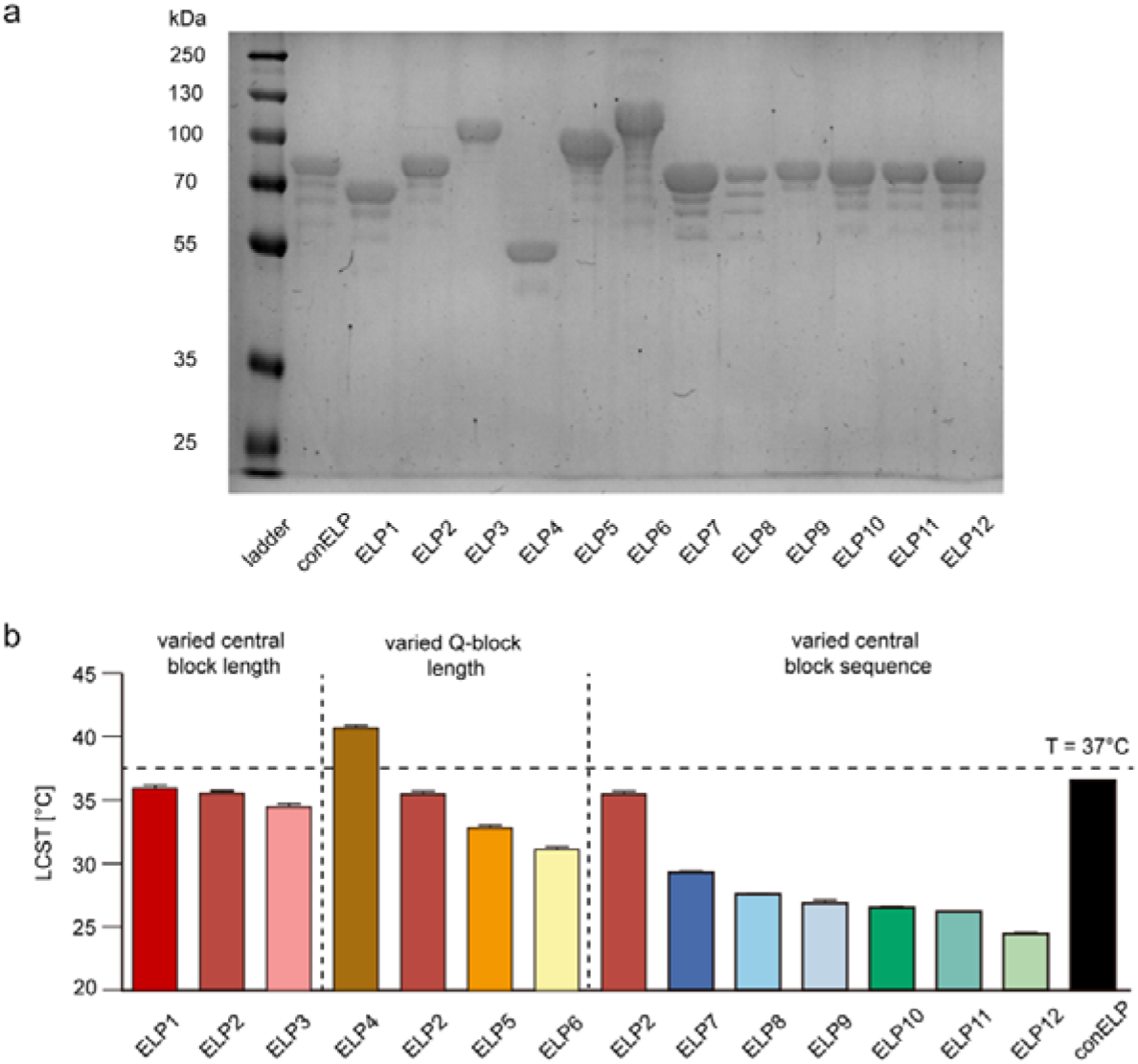
Molecular characterization and thermal phase behavior of Q-block-ELP constructs. SDS-PAGE analysis of Q-block-ELPs showing bands consistent with the expected molecular weights for the 12 Q-block-ELP constructs and non-crosslinkable control ELP (conELP). Variations in band mobility reflect systematic differences in central block length, Q-block valency, and guest residue composition across the ELP library. (b) LCST values for each ELP were determined by a turbidity assay at 30 µM. Most Q-block-ELPs exhibited LCSTs below physiological temperature (dashed line at 37L°C), indicating their capacity to undergo phase separation and form coacervates *in vivo*. Constructs with increased hydrophobicity or Q-block valency demonstrated reduced LCSTs.

### Influence of Q-block-ELPs on Fibrin Mechanics

Fibrin provides the primary structural framework that gives blood clots their mechanical strength, which is essential for effective hemostasis, vessel occlusion, and clot stability.^30^ The mechanical properties of the fibrin network not only influence bleeding control but also affect a variety of biological processes, including cell adhesion, migration, proliferation, wound healing, angiogenesis, and inflammation.^31^ The fibrin network is characterized by its high elasticity and ability to resist deformation, both of which are essential for maintaining clot integrity under the dynamic shear and mechanical stresses of blood circulation.^32^ Therefore, we evaluated how different ELP constructs influence the stiffness of fibrin hydrogels by measuring the storage modulus (G′) under physiological conditions using low-strain oscillatory shear rheology, which reliably reports on clot stiffness. We prepared fibrin gels by mixing 3.0 mg/mL fibrinogen with 30 μM Q-block-ELP, 20 mM CaCl_2_, and 0.2 U/mL thrombin at 37 °C. We performed rheological measurements at a strain amplitude of 1% over a frequency range of 0.1–3 Hz (γ = 1%). The maximum G′ reached within 3600 s was used to define the steady-state stiffness of each hybrid ELP/fibrin gel.^24^

As shown in Figure 3a, when the number of Q-blocks (i.e., cross-linkable glutamine residues) is held constant, increasing the length of the central ELP block leads to a modest reduction in fibrin network stiffness. This likely reflects greater flexibility within the ELP linker region of the gel. As the central block lengthens, the increased segment length reduces the tension held in the linker regions, thereby lowering the mechanical stiffness of the network.

**Figure 3.**
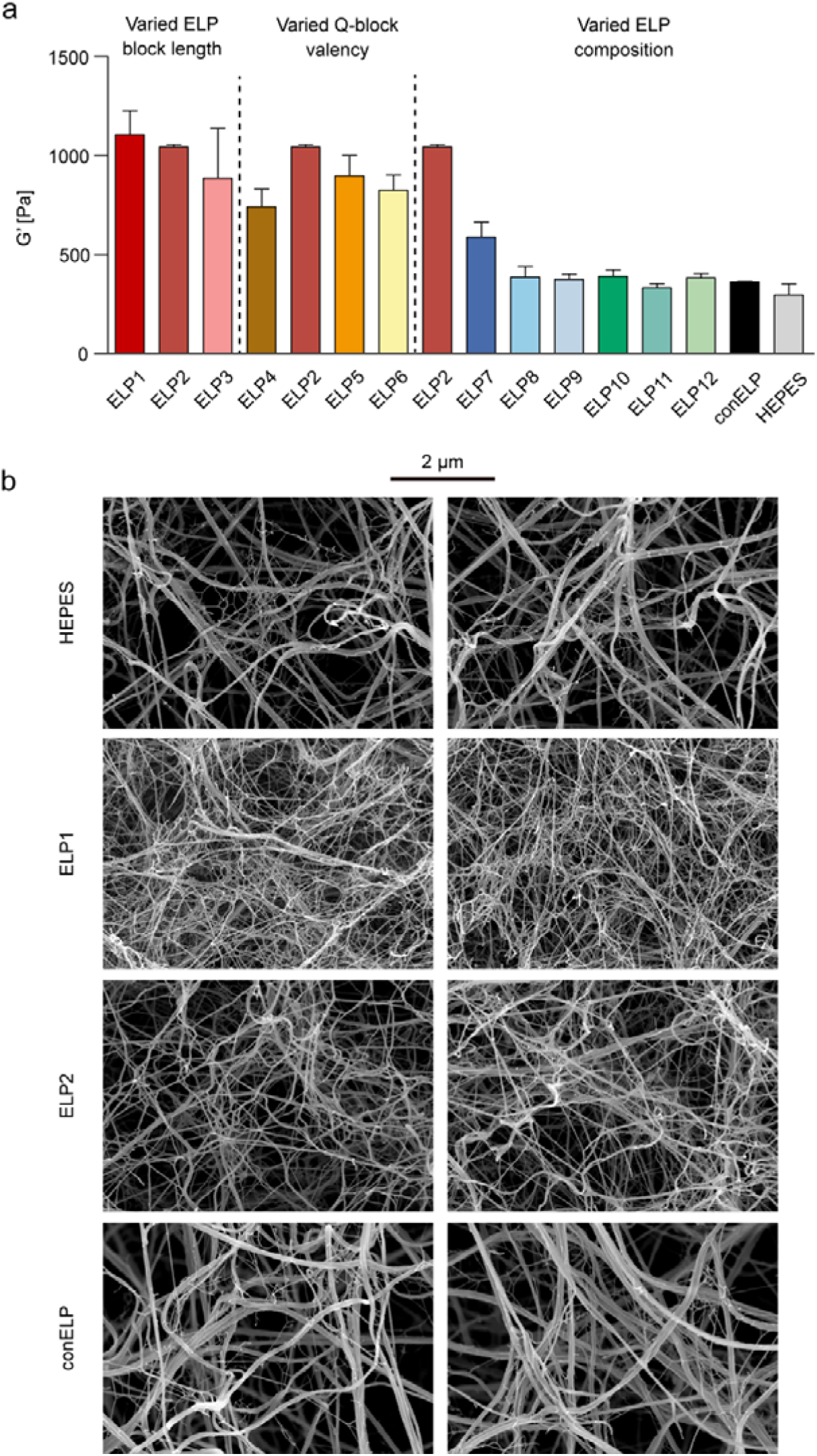
Q-block-ELPs enhance fibrin network stiffness and alter microarchitecture. (a) Oscillatory shear rheology was used to measure the storage modulus (G′) of fibrin hydrogels polymerized in the presence of different Q-block-ELP constructs. Several constructs (e.g., ELP1, ELP2, and ELP5) significantly increased fibrin gel stiffness relative to buffer control (HEPES) or the inactive conELP variant, indicating that both cross-linking valency and polymer architecture influence clot mechanics. (b) Scanning electron microscopy (SEM) images of fibrin clots show structural differences at the nanoscale. Fibrin treated with ELP1 and ELP2 formed denser networks composed of thinner and more highly interconnected fibers compared to conELP or HEPES controls, which showed a more porous structure.

In contrast, when the central block length is fixed, increasing the number of Q-blocks enhances the number of available glutamine cross-links, which initially results in greater fibrin cross-linking and increased network stiffness. This effect is observed when comparing constructs from ELP4 to ELP2. However, beyond a certain point (ELP2 to ELP5/ELP6), further increases in Q-block valency no longer yield proportional gains in stiffness. This plateau suggests that cross-linking capacity may become saturated, while continued increases in overall ELP length once again reduce network stiffness.

In constructs where guest residue hydrophobicity was increased, fibrin stiffness decreased further. This trend paralleled the LCST behavior of the constructs, suggesting that increased hydrophobicity and tighter aggregation through phase separation may limit accessibility of the Q-block glutamines for reaction with Lysine residues, thereby reducing effective cross-linking and network stiffness. Taken together, these findings indicate that optimal tuning of LCST and mechanical reinforcement requires careful balance of total ELP length, Q-block valency, and hydrophobicity.

### SEM Imaging Reveals ELP-Mediated Alterations in Fibrin Network Architecture

To evaluate the effects of ELP incorporation on fibrin network architecture at the ultrastructural level, we performed scanning electron microscopy (SEM) on fibrin clots polymerized at 37°C. Clots were formed by combining 1.5 mg/mL fibrinogen, 20 mM CaCl₂, 0.2 U/mL thrombin, and either buffer control (HEPES), 30 μM conELP, 30 μM ELP1, or 30 μM ELP2 (**Figure 3b**). In both the HEPES and conELP groups, fibrin networks displayed a porous structure composed of thick fibers with relatively low cross-linking density. In contrast, fibrin clots formed in the presence of ELP1 or ELP2 exhibited a denser network of thinner fibers, consistent with increased cross-linking within the fibrin scaffold. These findings provide morphological evidence that Q-block-ELPs are directly incorporated into the fibrin network via FXIIIa-mediated cross-linking. The microstructure observed with conELP, which lacks functional glutamine residues, was very similar to the HEPES control, confirming that the observed structural changes are specific to covalent incorporation of Q-block ELPs rather than nonspecific interactions with the ELP. The nanoscale organization of fibrin networks is known to contribute to mechanical properties of clots as well as their susceptibility to enzymatic degradation (i.e. fibrinolysis).^30, 33^ Collectively, these data suggest that Q-block ELPs may enhance hemostatic function by simultaneously increasing clot stiffness and resistance to enzymatic degradation.

### Covalent Incorporation of Q-block-ELPs into the Fibrin Network

To directly assess whether Q-block-ELPs are covalently incorporated into fibrin, we evaluated their cross-linking with fibrinogen in the presence of thrombin and CaCl₂ using SDS-PAGE (**Figure 4a**). In the absence of cross-linking, three distinct bands corresponding to the α, β, and γ chains of fibrinogen were visible (lane 2). Upon addition of thrombin and CaCl₂, fibrin was able to polymerize and the fibrinogen monomer bands disappeared (lane 7). When Q-block-bearing ELP1 or ELP2 were added to the fibrinogen mixture together with thrombin and CaCl₂, we observed a substantial reduction in the band intensity corresponding to the ELP monomer for both ELP1 and ELP2 (lanes 4 and 8; lanes 5 and 9), indicating covalent incorporation of the ELPs into the fibrin network. In contrast, the conELP negative control which lacks the Q-residues necessary for FXIIIa-mediated cross-linking showed no reduction in ELP monomer band intensity (lanes 6 and 10), confirming the specificity of cross-linking through the Q-residue within the Q-block sequences. To further demonstrate the cross-linking of ELP with fibrinogen, ELP1, ELP2, and the negative control conELP (10 µM) were all incubated with 1 µM of FITC-labeled fibrinogen γ 400-411 peptide (peptide sequences in the fibrinogen γ chain containing cross-linking sites with ELP) and factor XIIIa/CaCl₂. We first determined the corresponding band positions of ELP protein in the SDS-PAGE gel using Coomassie Blue staining in Figure S1A, and then determined the position of fibrinogen γ 400-411 using FITC fluorescence. We found that fibrinogen γ 400-411 showed fluorescence signals at the corresponding band positions of ELP1 and ELP2 proteins, while no fluorescence signal was found at the position of the control group conELP. The peptide becomes part of a covalent ELP-peptide conjugate, which demonstrates the chemical cross-linking of ELP with fibrinogen.

**Figure 4.**
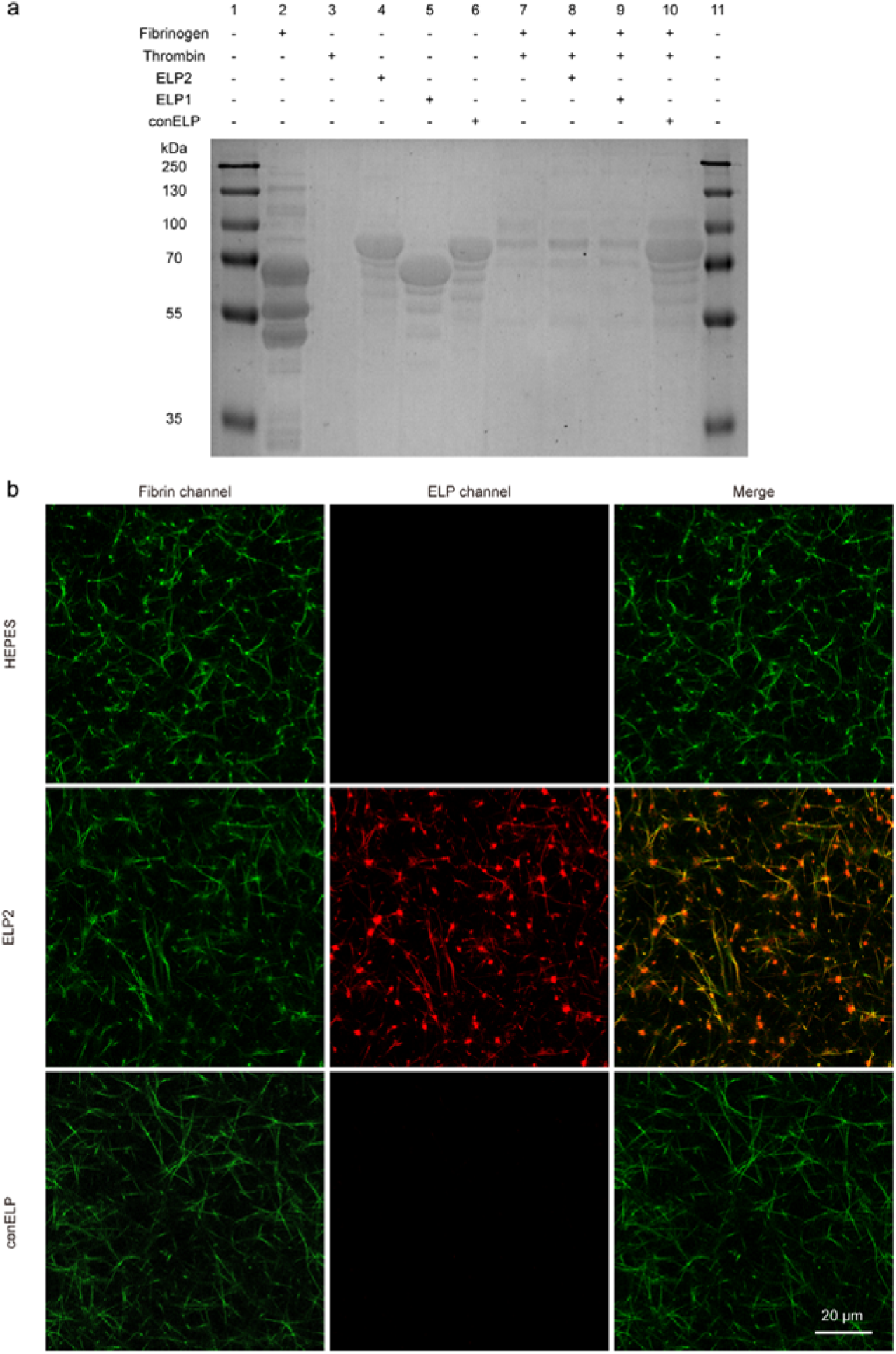
Q-block-ELPs are covalently integrated into fibrin networks via FXIIIa activity. (a) SDS-PAGE analysis of fibrinogen cross-linking reactions in the presence or absence of thrombin and various ELP constructs. In the presence of thrombin and Ca²⁺ (lanes 8–9), ELP1 and ELP2 bands are depleted compared to non-crosslinked controls (lanes 4–5) due to covalent bonding to fibrin. The conELP lacking Q-residues remains uncrosslinked (lane 10 vs. lane 6). (b) Two-color confocal fluorescence microscopy of fibrin clots formed with Alexa Fluor 488-labeled fibrinogen (green) and Alexa Fluor 594-labeled ELP2 or conELP (red). ELP2 shows strong co-localization with fibrin fibers. Minimal signal is observed in the ELP channel for clots formed with conELP or buffer alone. The punctate structure of red fluorescence is consistent with coacervate formation by ELP2 above its LCST.

To further verify chemical incorporation of ELP into the fibrin network, we fluorescently labeled ELP2 and conELP at their N-termini with Alexa Fluor 594 (red channel), and polymerized them with Alexa Fluor 488-labeled fibrinogen (green channel). Confocal fluorescence microscopy demonstrated fibrin network formation in all samples (HEPES, ELP2, and conELP groups) in the green channel (**Figure 4b**). However, strong red fluorescence signals were observed only in the ELP2-containing clots, where ELP2 co-localized with fibrin fibers, confirming its covalent incorporation into the fibrin matrix. Minimal red signal was detected in the HEPES and conELP groups, consistent with the absence of cross-linking. In addition, discrete ELP-rich coacervates were visible as bright red puncta in the ELP2 group, reflecting phase-separated domains that likely serve as additional cross-linking hubs within the fibrin scaffold. These findings further support the successful and specific incorporation of Q-block ELPs into fibrin networks above their LCST and help explain the structural basis for their ability to enhance fibrin mechanical properties.

### Platelet Depletion Model Using Anti-CD41 Antibody

Current therapeutic options for thrombocytopenia include plasma, platelet, coagulation factor, and fibrinogen concentrate transfusions, as well as antifibrinolytic agents such as tranexamic acid (TXA). However, plasma-derived products remain costly, require specialized storage, and carry risks of viral transmission and immunogenicity. Although TXA has been investigated in clinical studies for its potential to reduce bleeding or minimize the need for plasma or platelet transfusions in thrombocytopenic patients, results to date have been modest, and antifibrinolytic therapy alone has not proven sufficient to manage bleeding risk in this setting.^34^ As such, there is a strong rationale for developing recombinant protein-based therapeutics that reinforce fibrin network stability independent of platelet function or plasma-derived products. Such an approach may offer both mechanistic advantages in promoting clot stability and practical benefits in terms of safety, reliability, and scalability for clinical use.

To evaluate the procoagulant effects of ELPs under thrombocytopenic conditions, we first established an *in vivo* mouse model of thrombocytopenia using an anti-CD41 antibody. The anti-CD41 antibody binds to the CD41 receptor (glycoprotein GPIIb) on the platelet surface, marking platelets for clearance by the RES via macrophage-mediated phagocytosis. Mice were administered a single intraperitoneal (i.p.) injection of Ultra-LEAF™ Purified Anti-Mouse CD41 Antibody (Clone MWReg 30, BioLegend) at doses of 1 mg/kg, 0.5 mg/kg, 0.1 mg/kg, or vehicle control (0 mg/kg). To monitor platelet depletion, serial blood samples (20 μL) were collected from the tail vein at baseline and at 1, 3, 6, and 24 hours post-injection, diluted in 20 μL of 12 mM EDTA, and analyzed by flow cytometry. As shown in **Figure S2**, administration of 1 mg/kg anti-CD41 resulted in rapid platelet depletion to approximately 23% of baseline (severe thrombocytopenia)^35^, which remained stable over 24 hours. Doses of 0.5 mg/kg and 0.1 mg/kg yielded platelet reductions to approximately 43% (moderate severity) and 60% (moderate but close to mild thrombocytopenia) of baseline, respectively, with stable counts maintained through 24 hours.

### Q-Block-ELPs Reduce Bleeding in a Thrombocytopenic Mouse Tail-transection Model

To assess the hemostatic effects of ELPs in thrombocytopenic mice, we utilized a standard tail transection bleeding model to quantify blood loss following vascular injury. Mice were first rendered thrombocytopenic using 0.5 mg/kg of anti-CD41 antibody, and experiments were performed 24 hours post-injection, once platelet counts had stabilized at target levels (approximately 43% of baseline levels, corresponding to moderate Thrombocytopenia). Animals were anesthetized with isoflurane, and a 1 mm segment was excised from the distal tip of the tail to induce bleeding. Immediately following tail transection, mice received an intraperitoneal injection of ELP2 at doses of 0, 50, or 150 mg/kg. Intraperitoneal injections were chosen as they mimic continuous injections the closest, sustaining a relevant drug concentration in the blood for the period of the experiment (Figure S3). The tails were then immersed in pre-warmed (37°C) saline, and blood loss was collected in two consecutive 30-minute intervals, for a total of 60 minutes. At the conclusion of the bleeding assay, mice were euthanized by CO_2_ asphyxiation. Total blood loss was quantified by measuring hemoglobin concentration spectrophotometrically and calculating volume loss based on a hemoglobin standard curve.

As shown in **Figure 5b** (female mice) and **Figure 5f** (male mice), there was substantial variability in blood loss among individual animals within each group, which is consistent with previously reported variability inherent to the tail transection model.^36^ Although differences between groups did not reach statistical significance, likely due to sample size limitations, the data showed a clear dose-dependent trend. In female mice, average blood loss decreased from 461.9 ± 61.0 µL in controls to 404.5 ± 80.8 µL at 50 mg/kg and 296.5 ± 70.1 µL at 150 mg/kg ELP2 dosing. In male mice, blood loss decreased from 330.6 ± 51.3 µL in controls to 328.6 ± 54.9 µL at 50 mg/kg and 197.8 ± 68.1 µL at 150 mg/kg, with maximal blood loss reduction approaching ∼60% of control values. Additionally, in a subset of mice receiving the higher ELP dose, complete cessation of bleeding within 30 minutes was observed, which was not seen in control animals. These results suggest that treatment with Q-block-ELPs may promote hemostasis in a setting of moderate to severe thrombocytopenia. As shown in Figure S8, tranexamic acid (TA), a widely used antifibrinolytic drug, at doses (100 mg/kg, 4 repeated intraperitoneal administration) was tested as a clinical comparator. Interestingly, in our thrombocytopenic tail-clip model, TA infusion did not significantly reduce blood loss compared with the HEPES control, likely due to the distinct mechanism of TA, which primarily inhibits fibrinolysis rather than promoting clot formation under platelet-deficient conditions.

**Figure 5.**
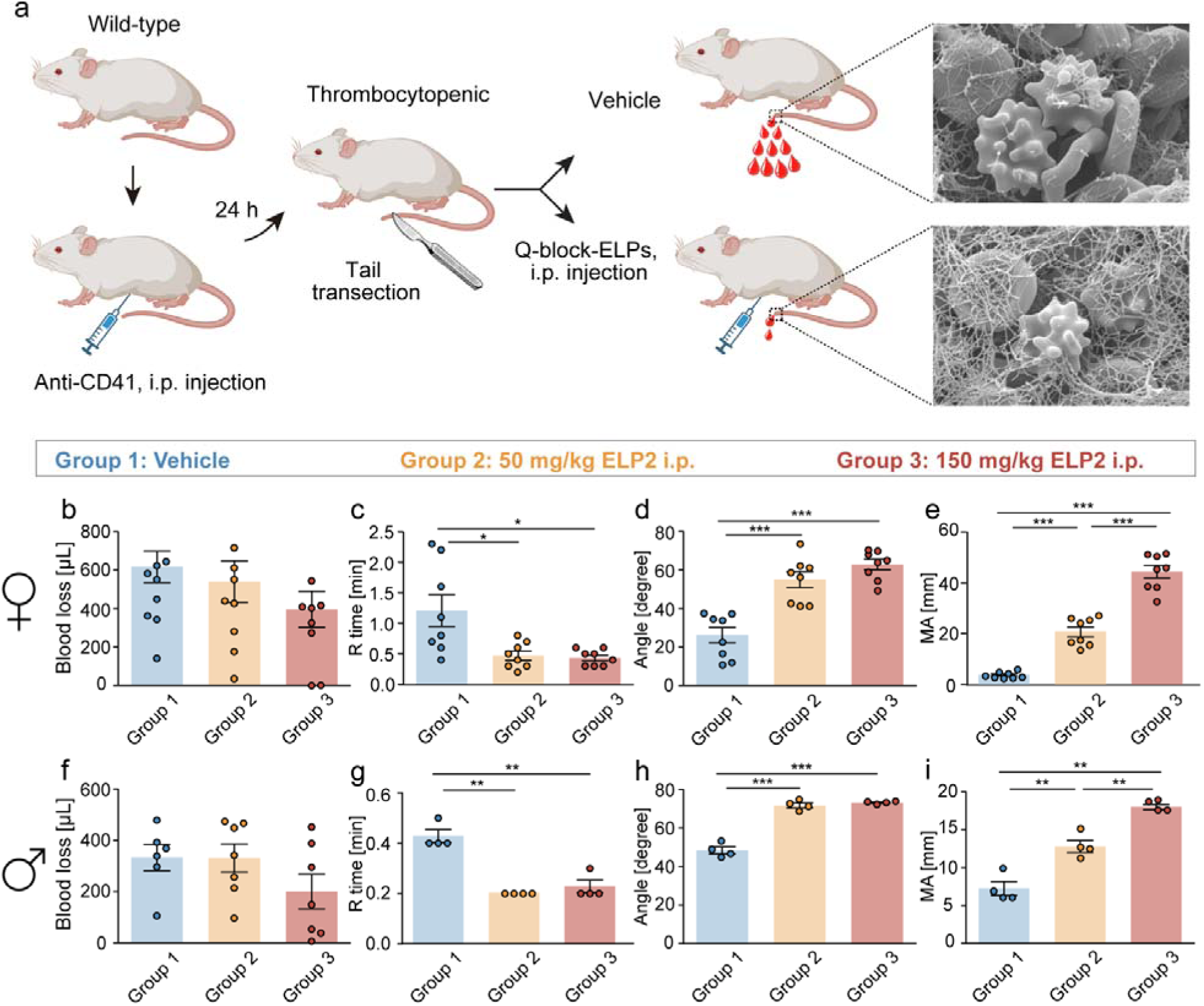
ELP administration reduced blood loss amount and increased kinetics and mechanical stability of blood clots from platelets depleted mice. a. Schematics of ELP i.p. injection in platelets depleted mice with tail transection bleeding model. ELP coacervates function as crosslinkers strengthening the fibrin network around platelets and red blood cells, which further stabilize mechanical property of blood clots in wound sites. b. Blood loss amounts of female mice in different groups (Group 1, 0 mg/kg of ELP i.p. injection, Group 2, 50 mg/kg of ELP i.p. injection, Group 3, 150 mg/kg of ELP i.p. injection) from tail transection bleeding model. c. Clotting onset time (R time), d. α-angle, and e. maximum amplitude (MA) of blood clots from female mice at different groups. f. Blood loss amounts of male mice in different groups from tail transection bleeding model. g. Clotting onset time (R time), h. α-angle, and i. maximum amplitude (MA) of blood clots from male mice at different groups.

### Biodistribution

To assess the in vivo distribution of ELP after injection, we performed whole-animal fluorescence imaging using ELP2 labeled with Cyanine 7.5 NHS ester. Mice were intraperitoneally injected with labeled ELP2 at 150 mg/kg, and fluorescence was monitored at multiple time points. As shown in **Figure S5b**, signal intensity increased rapidly within the first 3 hours, followed by a slower rise and eventual plateau through 48 hours, consistent with gradual tissue diffusion (**Figure S5a**). Limited by resolution, we could only determine that the ELP was localized to the abdominal cavity. To better define the tissue distribution, we harvested major organs at the terminal timepoint and performed ex vivo fluorescence imaging (**Figure S6**). The fluorescence intensity was highest in the intestines, followed by the stomach and liver, with small amounts detected in the pancreas and kidney.

We next assessed the pharmacokinetic profile of ELP2 following intraperitoneal administration by tracking fluorescently labeled ELP2 in the circulation over time. As shown in **Figure S3**, plasma concentrations of cyanine 7.5-labeled ELP2 were <5 μg/mL within the first 30 minutes post-injection. By 1 hour, concentrations had increased to 12.72 ± 6.77 μg/mL, reaching a peak level of 88.32 ± 50.50 μg/mL at 6 hours. Plasma concentrations subsequently declined to 28.40 ± 13.72 μg/mL by 24 hours post-injection. Minimal background fluorescence was detected in control animals receiving HEPES injection alone. However, this did not influence the pharmacokinetic trends observed in the ELP-treated animals.

To evaluate potential immunogenicity, we measured circulating total IgG and IgM levels 7 days after ELP administration using ELISA. This experiment was designed to provide a preliminary screen for acute immune activation. In mice, IgM responses peak around day 7, and early IgG can begin to appear. Therefore, we selected day 7 as a representative time point to evaluate whether a single administration triggers an acute antibody response. As shown in **Figure S4**, we did not observe any elevation of circulating IgM or IgG compared with controls. As ELP-specific antibodies are not commercially available currently, we incorporated published data on LPS-induced polyclonal activation as a benchmark. LPS is a potent activator of the immune system and has been repeatedly shown to induce a 5–10-fold increase in serum IgM within 48–96 hours in C57BL/6 mice^37^. This reference range is now indicated in **Figure S4** to illustrate the expected magnitude of an immune activation response. Relevant studies primarily investigated LPS-induced IgG subclass responses^38^. Since our assay quantifies total IgG, not IgG subclasses, existing LPS-based data cannot be directly compared. Therefore, we did not include LPS as a positive control in our IgG data. Normal physiological ranges of IgG and IgM are now shown as a gray band. To assist readers who may not be familiar with baseline immunoglobulin values, we have added gray reference bands representing the reported normal serum ranges for C57BL/6 mice: IgM 0.1–0.6 mg/mL and IgG 0.8–2.0 mg/mL^39^. As shown in the figure, our measured IgM and IgG fall within or near the lower end of the reported physiological range, confirming that no acute systemic immune activation was induced by the treatment. We acknowledge that this limited dataset cannot address delayed or cumulative immunogenicity.

### ELP-Mediated Modulation of Clot Kinetics and Stiffness by Thromboelastography

Thromboelastography (TEG) is a clinical method for evaluating the viscoelastic properties of whole blood during coagulation, providing real-time assessment of clot formation kinetics and mechanical strength.^40^ During TEG measurements, clot formation is monitored by measuring the displacement of a suspended pin within a rotating cup containing the blood sample. To characterize clotting dynamics, we analyzed standard TEG parameters. Reaction time (R time) reflects the time to initial fibrin formation, defined as the time required for the pin displacement to reach a 2 mm amplitude. The α-angle represents the steepness of the amplitude vs. time curve, and reports the rate of fibrin polymerization. Finally, maximum amplitude (MA) quantifies the mechanical strength of the clot, corresponding to its maximal viscoelastic resistance to deformation. To assess the effects of Q-block-ELPs on clot mechanics under thrombocytopenic conditions, blood was collected by cardiac puncture immediately following the tail transection bleeding experiments and measured by TEG.

As shown in **Figures 5c and 5g**, the significantly shortened R time in both female and male thrombocytopenic mice following administration of Q-block-ELPs indicated quicker initiation of clotting. In female mice, the R time was reduced from 1.2 minutes to 0.4 - 0.5 minutes following treatment with Q-block-ELPs. In males, R time decreased from 0.4 minutes to approximately 0.2 minutes, representing a 2–3-fold decrease in clot initiation time. The α-angle increased following Q-block-ELP treatment (**Figures 5d and 5h**). In female mice, the angle rose from 26° in controls to 55° with low-dose (50 mg/kg) Q-block-ELPs, and 63° with high-dose (150 mg/kg) Q-block-ELP treatment. In male mice, the angle increased from 48° to 71° and 73°, respectively, for low-dose and high-dose Q-block-ELP treated groups, indicating an up to 2.4-fold increase in clot formation rate. The MA increased in response to Q-block-ELP treatment (**Figures 5e and 5i**). In females, the MA rose from 4 mm to 21 mm with low-dose ELP, and 41 mm with high-dose ELP. In males, MA values increased from 7 mm to 13 mm and 18 mm in low-dose and high-dose groups, respectively, indicating an increase in clot strength of up to 10-fold. Sex-based differences in baseline values and response magnitude could reflect inherent sex-based variations in coagulability, clotting, and platelet biology. It has been demonstrated that males generally exhibit higher baseline platelet counts and enhanced aggregation potential, whereas females may display stronger fibrinolytic activity and differential hormone-related regulation of platelet activation^41^. Such sex-dependent coagulation differences could underlie the varying responses to ELP treatment. Together, the TEG results show that Q-block-ELPs accelerate clot formation and improve clot stiffness in thrombocytopenic mice. We also evaluated the effects of ELP on the kinetics and mechanical stability of blood clots in human with normal platelet counts. As shown in **Figure S7**, we analyzed coagulation parameters and clot formation profiles using fresh human blood samples with normal platelet counts. The results demonstrated that ELPs, at concentrations up to 30 μM, did not accelerate thrombus formation (R time) nor increase clot stiffness (MA value). The TEG parameters remained within physiological ranges, indicating that ELPs do not enhance systemic coagulability under normal hematologic conditions.

To investigate a structural basis for these viscoelastic changes, we performed SEM imaging of residual TEG samples (**Figure 6**). In untreated thrombocytopenic blood, minimal fibrin networks were observed around platelets and red blood cells. In contrast, samples treated with Q-block-ELPs, particularly at high dosing exhibited dense, well-organized fibrin networks surrounding cellular elements. These findings support the notion that Q-block-ELPs alter clot architecture at the nanoscale, translating into improved mechanical integrity and hemostatic function.

**Figure 6.**
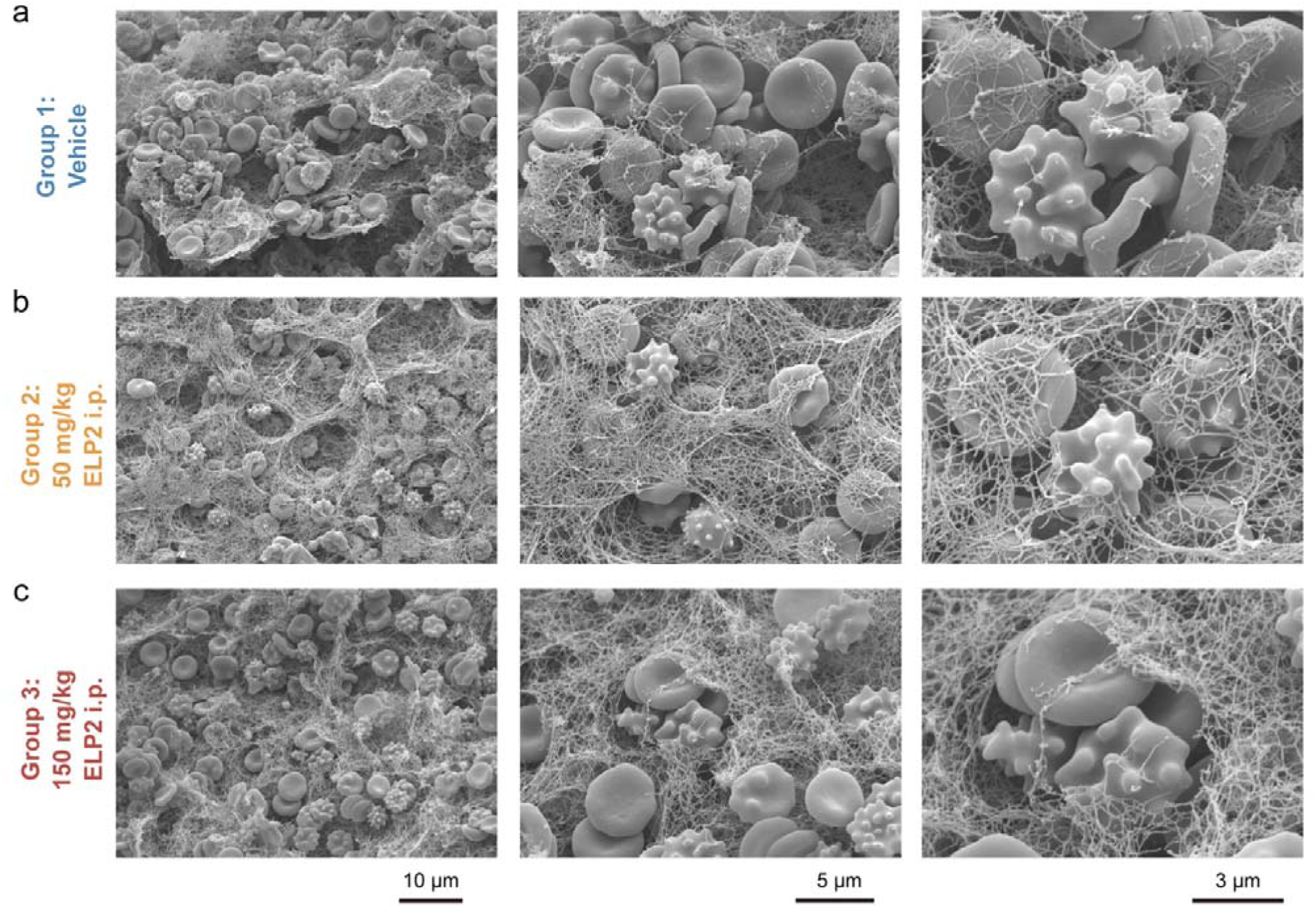
Ultrastructural analysis of clot morphology in thrombocytopenic mice treated with Q-block-ELPs. Scanning electron microscopy (SEM) images of clotted blood samples collected from thrombocytopenic mice following tail transection and treatment with either vehicle (a), 50 mg/kg ELP2 (b), or 150 mg/kg ELP2 (c). In vehicle controls (Group 1), the fibrin clot network is sparse with few platelet aggregates. In groups treated with Q-block-ELPs (Groups 2 and 3), denser fiber networks are observed, with integration of blood cells and platelets within the clot matrix. Samples from male and female mouse treatment groups were pooled prior to analysis.

## Conclusion

We report a protein-engineering strategy for enhancing blood clot mechanics in the setting of thrombocytopenia by combining an enzyme-responsive design with phase-separating protein polymers. By genetically encoding glutamine-containing Q-blocks at the termini of ELPs, we created constructs that serve as selective substrates for activated FXIIIa that are covalently bonded into nascent fibrin networks at sites of active clot formation *in vivo.* Above their LCST, these Q-block-ELPs undergo phase separation into protein-rich coacervates, which further reinforce clot structure through localized and physical aggregation. Across a rationally design library of 12 Q-block-ELPs with tunable hydrophobicity, length, and Q-block valency, we identified sequence-function relations that govern both phase behavior and mechanical performance of fibrin gels. In a mouse model of thrombocytopenia, systemic administration of Q-block-ELPs significantly accelerated clot formation, and increased clot stiffness.

This work demonstrates a modular framework for engineering recombinant protein materials that couple enzyme-responsiveness, supramolecular self-assembly, and mechanical reinforcement within the native clotting environment. By relying on the spatial and temporal control of FXIII-activation at sites undergoing active clotting, our Q-block-ELPs achieve clot-specific integration without crosslinking in healthy vasculature, helping to mitigate concerns about off-target thrombosis. Overall, these findings illustrate how programmable protein polymers can be leveraged to modulate clot architecture and mechanical function *in vivo*, and offer a blueprint for the development of next-generation biomaterials for bleeding control and hemostatic support.

## Author Contribution Statement

Y.S. designed and produced ELP constructs, conducted *in vitro* experiments, performed animal studies, analyzed data and wrote the manuscript. A.B. performed animal studies, analyzed data, and co-wrote the manuscript. I.U. provided key materials. S.K. and R.V. contributed to *in vitro* characterization and technical support, respectively. M.A.N. conceived of the project, secured funding, supervised the project, and co-wrote the manuscript with input from all authors.

## Acknowledgement

This work was supported by the University of Basel, ETH Zurich, and the European Research Council (Starting Grant; MMA - 715207). The authors would like to thank Michael Pfeffer for support in MS measurements and analysis, and Eva Bieler at the Nano Imaging Lab, University of Basel for her assistance with acquiring SEM images.

