## Supplementary Information for "Enzyme-responsive Hemostatic Elastin-like Polypeptides for Fibrin Stabilization and Coagulation Restoration in Thrombocytopenia"

### Methods and Materials

All chemicals were purchased from Sigma-Aldrich unless otherwise stated. Plasmids were synthesized by GeneArt (Thermo Scientific). Human fibrinogen (FIB 3, plasminogen, fibronectin, and von Willebrand Factor depleted), thrombin, and FXIIIa were purchased from Milan Analytica, AG (Rheinfelden, Switzerland).

*Sequences of ELP constructs*

ELP1

MGHGVGVPGVGVPGAGVPGVGVPGVGVPGVGVPGVGVPGVGVPGVGVPGVGVPGVGVPGEGVPGAGSGDQMM
LPWPAVALSGVPGVGVPGVGVPGAGVPGVGVPGVGVPGVGVPGVGVPGVGVPGVGVPGVGVPGVGVPGEGVPGA
GSGDQMMLPWPAVALSGVPGVGVPGVGVPGAGVPGVGVPGVGVPGVGVPGVGVPGVGVPGVGVPGVGVPGVGV
PGEGVPGAGSGDQMMLPWPAVALSGVPGVGVPGVGVPGAGVPGVGVPGVGVPGVGVPGVGVPGVGVPGVGVPG
VGVPGVGVPGEGVPGAGSGDQMMLPWPAVALSGVPGVGVPGVGVPGAGVPGVGVPGVGVPGVGVPGVGVPGVG
VPGVGVPGVGVPGVGVPGEGVPGAGVPGVGVPGVGVPGAGVPGVGVPGVGVPGVGVPGVGVPGVGVPGVGVPG
VGVPGVGVPGEGVPGAGVPGVGVPGVGVPGAGVPGVGVPGVGVPGVGVPGVGVPGVGVPGVGVPGVGVPGVGV
PGEGVPGAGSGDQMMLPWPAVALSGVPGVGVPGVGVPGAGVPGVGVPGVGVPGVGVPGVGVPGVGVPGVGVPG
VGVPGVGVPGEGVPGAGSGDQMMLPWPAVALSGVPGVGVPGVGVPGAGVPGVGVPGVGVPGVGVPGVGVPGVG
VPGVGVPGVGVPGVGVPGEGVPGAGSGDQMMLPWPAVALSGVPGVGVPGVGVPGAGVPGVGVPGVGVPGVGVP
GVGVPGVGVPGVGVPGVGVPGVGVPGEGVPGAGSGDQMMLPWPAVALSGVPGWLDSLEFIA*

ELP2

MGHGVGVPGVGVPGAGVPGVGVPGVGVPGVGVPGVGVPGVGVPGVGVPGVGVPGVGVPGEGVPGAGSGDQMM
LPWPAVALSGVPGVGVPGVGVPGAGVPGVGVPGVGVPGVGVPGVGVPGVGVPGVGVPGVGVPGVGVPGEGVPGA
GSGDQMMLPWPAVALSGVPGVGVPGVGVPGAGVPGVGVPGVGVPGVGVPGVGVPGVGVPGVGVPGVGVPGVGV
PGEGVPGAGSGDQMMLPWPAVALSGVPGVGVPGVGVPGAGVPGVGVPGVGVPGVGVPGVGVPGVGVPGVGVPG
VGVPGVGVPGEGVPGAGSGDQMMLPWPAVALSGVPGVGVPGVGVPGAGVPGVGVPGVGVPGVGVPGVGVPGVG
VPGVGVPGVGVPGVGVPGEGVPGAGVPGVGVPGVGVPGAGVPGVGVPGVGVPGVGVPGVGVPGVGVPGVGVPG
VGVPGVGVPGEGVPGAGVPGVGVPGVGVPGAGVPGVGVPGVGVPGVGVPGVGVPGVGVPGVGVPGVGVPGVGV
PGEGVPGAGVPGVGVPGVGVPGAGVPGVGVPGVGVPGVGVPGVGVPGVGVPGVGVPGVGVPGVGVPGEGVPGA
GVPGVGVPGVGVPGAGVPGVGVPGVGVPGVGVPGVGVPGVGVPGVGVPGVGVPGVGVPGEGVPGAGSGDQMM
LPWPAVALSGVPGVGVPGVGVPGAGVPGVGVPGVGVPGVGVPGVGVPGVGVPGVGVPGVGVPGVGVPGEGVPGA
GSGDQMMLPWPAVALSGVPGVGVPGVGVPGAGVPGVGVPGVGVPGVGVPGVGVPGVGVPGVGVPGVGVPGVGV
PGEGVPGAGSGDQMMLPWPAVALSGVPGVGVPGVGVPGAGVPGVGVPGVGVPGVGVPGVGVPGVGVPGVGVPG
VGVPGVGVPGEGVPGAGSGDQMMLPWPAVALSGVPGWLDSLEFIA*

ELP3

MGHGVGVPGVGVPGAGVPGVGVPGVGVPGVGVPGVGVPGVGVPGVGVPGVGVPGVGVPGEGVPGAGSGDQMM
LPWPAVALSGVPGVGVPGVGVPGAGVPGVGVPGVGVPGVGVPGVGVPGVGVPGVGVPGVGVPGVGVPGEGVPGA
GSGDQMMLPWPAVALSGVPGVGVPGVGVPGAGVPGVGVPGVGVPGVGVPGVGVPGVGVPGVGVPGVGVPGVGV
PGEGVPGAGSGDQMMLPWPAVALSGVPGVGVPGVGVPGAGVPGVGVPGVGVPGVGVPGVGVPGVGVPGVGVPG
VGVPGVGVPGEGVPGAGSGDQMMLPWPAVALSGVPGVGVPGVGVPGAGVPGVGVPGVGVPGVGVPGVGVPGVG
VPGVGVPGVGVPGVGVPGEGVPGAGVPGVGVPGVGVPGAGVPGVGVPGVGVPGVGVPGVGVPGVGVPGVGVPG
VGVPGVGVPGEGVPGAGVPGVGVPGVGVPGAGVPGVGVPGVGVPGVGVPGVGVPGVGVPGVGVPGVGVPGVGV
PGEGVPGAGVPGVGVPGVGVPGAGVPGVGVPGVGVPGVGVPGVGVPGVGVPGVGVPGVGVPGVGVPGEGVPGA

GVPGVGVPGVGVPGAGVPGVGVPGVGVPGVGVPGVGVPGVGVPGVGVPGVGVPGEGVPGAGVPGVGVP
GVGVPGAGVPGVGVPGVGVPGVGVPGVGVPGVGVPGVGVPGVGVPGEGVPGAGVPGVGVPGVGVPGAG
VPGVGVPGVGVPGVGVPGVGVPGVGVPGVGVPGVGVPGVGVPGEGVPGAGVPGVGVPGVGVPGAGVPGVGVPG
VGVPGVGVPGVGVPGVGVPGVGVPGVGVPGEGVPGAGVPGVGVPGVGVPGAGVPGVGVPGVGVPGVGV
PGVGVPGVGVPGVGVPGVGVPGEGVPGAGSGDQMMLPWPAVALSGVPGVGVPGVGVPGAGVPGVGVPG
VGVPGVGVPGVGVPGVGVPGVGVPGVGVPGEGVPGAGSGDQMMLPWPAVALSGVPGVGVPGVGVPGAG
VPGVGVPGVGVPGVGVPGVGVPGVGVPGVGVPGVGVPGEGVPGAGSGDQMMLPWPAVALSGVPGVGVP
GVGVPGAGVPGVGVPGVGVPGVGVPGVGVPGVGVPGVGVPGVGVPGEGVPGAGSGDQMMLPWPAVALS
GVPGWLDSLEFIA*

ELP4
MGHGVGVPGVGVPGAGVPGVGVPGVGVPGVGVPGVGVPGVGVPGVGVPGVGVPGEGVPGAGSGDQMM
LPWPAVALSGVPGVGVPGVGVPGAGVPGVGVPGVGVPGVGVPGVGVPGVGVPGVGVPGVGVPGEGVPGA
GSGDQMMLPWPAVALSGVPGVGVPGVGVPGAGVPGVGVPGVGVPGVGVPGVGVPGVGVPGVGVPGVGV
PGEGVPGAGVPGVGVPGVGVPGAGVPGVGVPGVGVPGVGVPGVGVPGVGVPGVGVPGVGVPGEGVPGA
GVPGVGVPGVGVPGAGVPGVGVPGVGVPGVGVPGVGVPGVGVPGVGVPGVGVPGEGVPGAGVPGVGVP
GVGVPGAGVPGVGVPGVGVPGVGVPGVGVPGVGVPGVGVPGVGVPGEGVPGAGVPGVGVPGVGVPGAG
VPGVGVPGVGVPGVGVPGVGVPGVGVPGVGVPGVGVPGVGVPGEGVPGAGSGDQMMLPWPAVALSGVPGVGVP
GVGVPGAGVPGVGVPGVGVPGVGVPGVGVPGVGVPGVGVPGVGVPGEGVPGAGSGDQMMLPWPAVALS
GVPGWLDSLEFIA*

ELP5
MGHGVGVPGVGVPGAGVPGVGVPGVGVPGVGVPGVGVPGVGVPGVGVPGVGVPGEGVPGAGSGDQMM
LPWPAVALSGVPGVGVPGVGVPGAGVPGVGVPGVGVPGVGVPGVGVPGVGVPGVGVPGVGVPGEGVPGA
GSGDQMMLPWPAVALSGVPGVGVPGVGVPGAGVPGVGVPGVGVPGVGVPGVGVPGVGVPGVGVPGVGV
PGEGVPGAGSGDQMMLPWPAVALSGVPGVGVPGVGVPGAGVPGVGVPGVGVPGVGVPGVGVPGVGVPG
VGVPGVGVPGEGVPGAGSGDQMMLPWPAVALSGVPGVGVPGVGVPGAGVPGVGVPGVGVPGVGVPGVG
VPGVGVPGVGVPGVGVPGEGVPGAGSGDQMMLPWPAVALSGVPGVGVPGVGVPGAGVPGVGVPGVGVP
GVGVPGVGVPGVGVPGVGVPGVGVPGEGVPGAGSGDQMMLPWPAVALSGVPGVGVPGVGVPGAGVPGV
GVPGVGVPGVGVPGVGVPGVGVPGVGVPGVGVPGEGVPGAGVPGVGVPGVGVPGAGVPGVGVPGVGVP
GVGVPGVGVPGVGVPGVGVPGVGVPGEGVPGAGVPGVGVPGVGVPGAGVPGVGVPGVGVPGVGVPGVG
VPGVGVPGVGVPGVGVPGEGVPGAGVPGVGVPGVGVPGAGVPGVGVPGVGVPGVGVPGVGVPGVGVPG
VGVPGVGVPGEGVPGAGVPGVGVPGVGVPGAGVPGVGVPGVGVPGVGVPGVGVPGVGVPGVGVPGVGV
PGEGVPGAGSGDQMMLPWPAVALSGVPGVGVPGVGVPGAGVPGVGVPGVGVPGVGVPGVGVPGVGVPG
VGVPGVGVPGEGVPGAGSGDQMMLPWPAVALSGVPGVGVPGVGVPGAGVPGVGVPGVGVPGVGVPGVG
VPGVGVPGVGVPGVGVPGEGVPGAGSGDQMMLPWPAVALSGVPGVGVPGVGVPGAGVPGVGVPGVGVP
GVGVPGVGVPGVGVPGVGVPGVGVPGEGVPGAGSGDQMMLPWPAVALSGVPGVGVPGVGVPGAGVPGV
GVPGVGVPGVGVPGVGVPGVGVPGVGVPGVGVPGEGVPGAGSGDQMMLPWPAVALSGVPGVGVPGVGV
PGAGVPGVGVPGVGVPGVGVPGVGVPGVGVPGVGVPGVGVPGEGVPGAGSGDQMMLPWPAVALSGVPG
WLDSLEFIA*

ELP6
MGHGVGVPGVGVPGAGVPGVGVPGVGVPGVGVPGVGVPGVGVPGVGVPGVGVPGEGVPGAGSGDQMM
LPWPAVALSGVPGVGVPGVGVPGAGVPGVGVPGVGVPGVGVPGVGVPGVGVPGVGVPGVGVPGEGVPGA

GSGDQMMLPWPAVALSGVPGVGVPGVGVPGAGVPGVGVPGVGVPGVGVPGVGVPGVGVPGVGVPGVGV
PGEGVPGAGSGDQMMLPWPAVALSGVPGVGVPGVGVPGAGVPGVGVPGVGVPGVGVPGVGVPGVGVPG
VGVPGVGVPGEGVPGAGSGDQMMLPWPAVALSGVPGVGVPGVGVPGAGVPGVGVPGVGVPGVGVPGVG
VPGVGVPGVGVPGVGVPGEGVPGAGSGDQMMLPWPAVALSGVPGVGVPGVGVPGAGVPGVGVPGVGVP
GVGVPGVGVPGVGVPGVGVPGVGVPGEGVPGAGSGDQMMLPWPAVALSGVPGVGVPGVGVPGAGVPGV
GVPGVGVPGVGVPGVGVPGVGVPGVGVPGVGVPGEGVPGAGSGDQMMLPWPAVALSGVPGVGVPGVGV
PGAGVPGVGVPGVGVPGVGVPGVGVPGVGVPGVGVPGVGVPGEGVPGAGSGDQMMLPWPAVALSGVPG
VGVPGVGVPGAGVPGVGVPGVGVPGVGVPGVGVPGVGVPGVGVPGVGVPGEGVPGAGVPGVGVPGVGV
PGAGVPGVGVPGVGVPGVGVPGVGVPGVGVPGVGVPGVGVPGEGVPGAGVPGVGVPGVGVPGAGVPGV
GVPGVGVPGVGVPGVGVPGVGVPGVGVPGVGVPGEGVPGAGVPGVGVPGVGVPGAGVPGVGVPGVGVP
GVGVPGVGVPGVGVPGVGVPGVGVPGEGVPGAGVPGVGVPGVGVPGAGVPGVGVPGVGVPGVGVPGVG
VPGVGVPGVGVPGVGVPGEGVPGAGSGDQMMLPWPAVALSGVPGVGVPGVGVPGAGVPGVGVPGVGVP
GVGVPGVGVPGVGVPGVGVPGVGVPGEGVPGAGSGDQMMLPWPAVALSGVPGVGVPGVGVPGAGVPGV
GVPGVGVPGVGVPGVGVPGVGVPGVGVPGVGVPGEGVPGAGSGDQMMLPWPAVALSGVPGVGVPGVGV
PGAGVPGVGVPGVGVPGVGVPGVGVPGVGVPGVGVPGVGVPGEGVPGAGSGDQMMLPWPAVALSGVPG
VGVPGVGVPGAGVPGVGVPGVGVPGVGVPGVGVPGVGVPGVGVPGVGVPGEGVPGAGSGDQMMLPWP
AVALSGVPGVGVPGVGVPGAGVPGVGVPGVGVPGVGVPGVGVPGVGVPGVGVPGVGVPGEGVPGAGSG
DQMMLPWPAVALSGVPGVGVPGVGVPGAGVPGVGVPGVGVPGVGVPGVGVPGVGVPGVGVPGVGVPGE
GVPGAGSGDQMMLPWPAVALSGVPGVGVPGVGVPGAGVPGVGVPGVGVPGVGVPGVGVPGVGVPGVGV
PGVGVPGEGVPGAGSGDQMMLPWPAVALSGVPGWLDSLEFIA*

ELP7
MGHGVGVPGVGVPGAGVPGVGVPGVGVPGVGVPGVGVPGVGVPGVGVPGVGVPGEGVPGAGSGDQMM
LPWPAVALSGVPGVGVPGVGVPGAGVPGVGVPGVGVPGVGVPGVGVPGVGVPGVGVPGVGVPGEGVPGA
GSGDQMMLPWPAVALSGVPGVGVPGVGVPGAGVPGVGVPGVGVPGVGVPGVGVPGVGVPGVGVPGVGV
PGEGVPGAGSGDQMMLPWPAVALSGVPGVGVPGVGVPGAGVPGVGVPGVGVPGVGVPGVGVPGVGVPG
VGVPGVGVPGEGVPGAGSGDQMMLPWPAVALSGVPGVGVPGVGVPGVGVPGVGVPGYGVPGVGVPGVG
VPGVGVPGVGVPGVGVPGYGVPGVGVPGVGVPGVGVPGVGVPGYGVPGVGVPGVGVPGVGVPGVGVPG
VGVPGYGVPGVGVPGVGVPGVGVPGVGVPGYGVPGVGVPGVGVPGVGVPGVGVPGVGVPGYGVPGVGV
PGVGVPGVGVPGVGVPGYGVPGVGVPGVGVPGVGVPGVGVPGVGVPGYGVPGVGVPGVGVPGAGVPGV
GVPGVGVPGVGVPGVGVPGVGVPGVGVPGVGVPGEGVPGAGSGDQMMLPWPAVALSGVPGVGVPGVGV
PGAGVPGVGVPGVGVPGVGVPGVGVPGVGVPGVGVPGVGVPGEGVPGAGSGDQMMLPWPAVALSGVPG
VGVPGVGVPGAGVPGVGVPGVGVPGVGVPGVGVPGVGVPGVGVPGVGVPGEGVPGAGSGDQMMLPWP
AVALSGVPGVGVPGVGVPGAGVPGVGVPGVGVPGVGVPGVGVPGVGVPGVGVPGVGVPGEGVPGAGSG
DQMMLPWPAVALSGVPGWLDSLEFIA*

ELP8
MGHGVGVPGVGVPGAGVPGVGVPGVGVPGVGVPGVGVPGVGVPGVGVPGVGVPGEGVPGAGSGDQMM
LPWPAVALSGVPGVGVPGVGVPGAGVPGVGVPGVGVPGVGVPGVGVPGVGVPGVGVPGVGVPGEGVPGA
GSGDQMMLPWPAVALSGVPGVGVPGVGVPGAGVPGVGVPGVGVPGVGVPGVGVPGVGVPGVGVPGVGV
PGEGVPGAGSGDQMMLPWPAVALSGVPGVGVPGVGVPGAGVPGVGVPGVGVPGVGVPGVGVPGVGVPG
VGVPGVGVPGEGVPGAGSGDQMMLPWPAVALSGVPGVGVPGVGVPGYGVPGVGVPGYGVPGVGVPGYG
VPGVGVPGYGVPGVGVPGYGVPGVGVPGVGVPGYGVPGVGVPGYGVPGVGVPGYGVPGVGVPGYGVPG
VGVPGYGVPGVGVPGVGVPGYGVPGVGVPGYGVPGVGVPGYGVPGVGVPGYGVPGVGVPGYGVPGVGV

PGVGVPGYGVPGVGVPGYGVPGVGVPGYGVPGVGVPGYGVPGVGVPGYGVPGVGVPGVGVPGAGVPGV
GVPGVGVPGVGVPGVGVPGVGVPGVGVPGVGVPGEGVPGAGSGDQMMLPWPAVALSGVPGVGVPGVGV
PGAGVPGVGVPGVGVPGVGVPGVGVPGVGVPGVGVPGVGVPGEGVPGAGSGDQMMLPWPAVALSGVPG
VGVPGVGVPGAGVPGVGVPGVGVPGVGVPGVGVPGVGVPGVGVPGVGVPGEGVPGAGSGDQMMLPWP
AVALSGVPGVGVPGVGVPGAGVPGVGVPGVGVPGVGVPGVGVPGVGVPGVGVPGVGVPGEGVPGAGSG
DQMMLPWPAVALSGVPGWLDSLEFIA*

ELP9
MGHGVGVPGVGVPGAGVPGVGVPGVGVPGVGVPGVGVPGVGVPGVGVPGVGVPGEGVPGAGSGDQMM
LPWPAVALSGVPGVGVPGVGVPGAGVPGVGVPGVGVPGVGVPGVGVPGVGVPGVGVPGVGVPGEGVPGA
GSGDQMMLPWPAVALSGVPGVGVPGVGVPGAGVPGVGVPGVGVPGVGVPGVGVPGVGVPGVGVPGVGV
PGEGVPGAGSGDQMMLPWPAVALSGVPGVGVPGVGVPGAGVPGVGVPGVGVPGVGVPGVGVPGVGVPG
VGVPGVGVPGEGVPGAGSGDQMMLPWPAVALSGVPGVGVPGVGVPGVGVPGVGVPGAGVPGVGVPGVG
VPGVGVPGVGVPGVGVPGYGVPGAGVPGVGVPGVGVPGVGVPGVGVPGAGVPGVGVPGVGVPGVGVPG
VGVPGVGVPGYGVPGAGVPGVGVPGVGVPGVGVPGVGVPGAGVPGVGVPGVGVPGVGVPGVGVPGVGV
PGYGVPGAGVPGVGVPGVGVPGVGVPGVGVPGAGVPGVGVPGVGVPGVGVPGVGVPGVGVPGYGVPGA
GVPGVGVPGVGVPGAGVPGVGVPGVGVPGVGVPGVGVPGVGVPGVGVPGVGVPGEGVPGAGSGDQMM
LPWPAVALSGVPGVGVPGVGVPGAGVPGVGVPGVGVPGVGVPGVGVPGVGVPGVGVPGVGVPGEGVPGA
GSGDQMMLPWPAVALSGVPGVGVPGVGVPGAGVPGVGVPGVGVPGVGVPGVGVPGVGVPGVGVPGVGV
PGEGVPGAGSGDQMMLPWPAVALSGVPGVGVPGVGVPGAGVPGVGVPGVGVPGVGVPGVGVPGVGVPG
VGVPGVGVPGEGVPGAGSGDQMMLPWPAVALSGVPGWLDSLEFIA*

ELP10
MGHGVGVPGVGVPGAGVPGVGVPGVGVPGVGVPGVGVPGVGVPGVGVPGVGVPGEGVPGAGSGDQMM
LPWPAVALSGVPGVGVPGVGVPGAGVPGVGVPGVGVPGVGVPGVGVPGVGVPGVGVPGVGVPGEGVPGA
GSGDQMMLPWPAVALSGVPGVGVPGVGVPGAGVPGVGVPGVGVPGVGVPGVGVPGVGVPGVGVPGVGV
PGEGVPGAGSGDQMMLPWPAVALSGVPGVGVPGVGVPGAGVPGVGVPGVGVPGVGVPGVGVPGVGVPG
VGVPGVGVPGEGVPGAGSGDQMMLPWPAVALSGVPGVGVPGVGVPGVGVPGVGVPGYGVPGVGVPGVG
VPGVGVPGVGVPGVGVPGYGVPGAGVPGVGVPGVGVPGVGVPGVGVPGYGVPGVGVPGVGVPGVGVPG
VGVPGVGVPGYGVPGAGVPGVGVPGVGVPGVGVPGVGVPGYGVPGVGVPGVGVPGVGVPGVGVPGVGV
PGYGVPGAGVPGVGVPGVGVPGVGVPGVGVPGYGVPGVGVPGVGVPGVGVPGVGVPGVGVPGYGVPGA
GVPGVGVPGVGVPGAGVPGVGVPGVGVPGVGVPGVGVPGVGVPGVGVPGVGVPGEGVPGAGSGDQMM
LPWPAVALSGVPGVGVPGVGVPGAGVPGVGVPGVGVPGVGVPGVGVPGVGVPGVGVPGVGVPGEGVPGA
GSGDQMMLPWPAVALSGVPGVGVPGVGVPGAGVPGVGVPGVGVPGVGVPGVGVPGVGVPGVGVPGVGV
PGEGVPGAGSGDQMMLPWPAVALSGVPGVGVPGVGVPGAGVPGVGVPGVGVPGVGVPGVGVPGVGVPG
VGVPGVGVPGEGVPGAGSGDQMMLPWPAVALSGVPGWLDSLEFIA*

ELP11
MGHGVGVPGVGVPGAGVPGVGVPGVGVPGVGVPGVGVPGVGVPGVGVPGVGVPGEGVPGAGSGDQMM
LPWPAVALSGVPGVGVPGVGVPGAGVPGVGVPGVGVPGVGVPGVGVPGVGVPGVGVPGVGVPGEGVPGA
GSGDQMMLPWPAVALSGVPGVGVPGVGVPGAGVPGVGVPGVGVPGVGVPGVGVPGVGVPGVGVPGVGV
PGEGVPGAGSGDQMMLPWPAVALSGVPGVGVPGVGVPGAGVPGVGVPGVGVPGVGVPGVGVPGVGVPG
VGVPGVGVPGEGVPGAGSGDQMMLPWPAVALSGVPGVGVPGVGVPGVGVPGVGVPGYGVPGVGVPGVG
VPGVGVPGVGVPGVGVPGYGVPGYGVPGVGVPGVGVPGVGVPGVGVPGYGVPGVGVPGVGVPGVGVPG

VGVPGVGVPGYGVPGYGVPGVGVPGVGVPGVGVPGVGVPGYGVPGVGVPGVGVPGVGVPGVGVPGVGV
PGYGVPGYGVPGVGVPGVGVPGVGVPGVGVPGYGVPGVGVPGVGVPGVGVPGVGVPGVGVPGYGVPGY
GVPGVGVPGVGVPGAGVPGVGVPGVGVPGVGVPGVGVPGVGVPGVGVPGVGVPGEGVPGAGSGDQMM
LPWPAVALSGVPGVGVPGVGVPGAGVPGVGVPGVGVPGVGVPGVGVPGVGVPGVGVPGVGVPGEGVPGA
GSGDQMMLPWPAVALSGVPGVGVPGVGVPGAGVPGVGVPGVGVPGVGVPGVGVPGVGVPGVGVPGVGV
PGEGVPGAGSGDQMMLPWPAVALSGVPGVGVPGVGVPGAGVPGVGVPGVGVPGVGVPGVGVPGVGVPG
VGVPGVGVPGEGVPGAGSGDQMMLPWPAVALSGVPGWLDSLEFIA*

ELP12
MGHGVGVPGVGVPGAGVPGVGVPGVGVPGVGVPGVGVPGVGVPGVGVPGVGVPGEGVPGAGSGDQMM
LPWPAVALSGVPGVGVPGVGVPGAGVPGVGVPGVGVPGVGVPGVGVPGVGVPGVGVPGVGVPGEGVPGA
GSGDQMMLPWPAVALSGVPGVGVPGVGVPGAGVPGVGVPGVGVPGVGVPGVGVPGVGVPGVGVPGVGV
PGEGVPGAGSGDQMMLPWPAVALSGVPGVGVPGVGVPGAGVPGVGVPGVGVPGVGVPGVGVPGVGVPG
VGVPGVGVPGEGVPGAGSGDQMMLPWPAVALSGVPGVGVPGVGVPGVGVPGVGVPGYGVPGVGVPGVG
VPGVGVPGVGVPGVGVPGYGVPGEGVPGVGVPGVGVPGVGVPGVGVPGYGVPGVGVPGVGVPGVGVPG
VGVPGVGVPGYGVPGEGVPGVGVPGVGVPGVGVPGVGVPGYGVPGVGVPGVGVPGVGVPGVGVPGVGV
PGYGVPGEGVPGVGVPGVGVPGVGVPGVGVPGYGVPGVGVPGVGVPGVGVPGVGVPGVGVPGYGVPGE
GVPGVGVPGVGVPGAGVPGVGVPGVGVPGVGVPGVGVPGVGVPGVGVPGVGVPGEGVPGAGSGDQMM
LPWPAVALSGVPGVGVPGVGVPGAGVPGVGVPGVGVPGVGVPGVGVPGVGVPGVGVPGVGVPGEGVPGA
GSGDQMMLPWPAVALSGVPGVGVPGVGVPGAGVPGVGVPGVGVPGVGVPGVGVPGVGVPGVGVPGVGV
PGEGVPGAGSGDQMMLPWPAVALSGVPGVGVPGVGVPGAGVPGVGVPGVGVPGVGVPGVGVPGVGVPG
VGVPGVGVPGEGVPGAGSGDQMMLPWPAVALSGVPGWLDSLEFIA*

conELP
MGHGVGVPGVGVPGAGVPGVGVPGVGVPGVGVPGVGVPGVGVPGVGVPGVGVPGEGVPGAGSGDGMM
LPWPAVALSGVPGVGVPGVGVPGAGVPGVGVPGVGVPGVGVPGVGVPGVGVPGVGVPGVGVPGEGVPGA
GSGDGMMLPWPAVALSGVPGVGVPGVGVPGAGVPGVGVPGVGVPGVGVPGVGVPGVGVPGVGVPGVGV
PGEGVPGAGSGDGMMLPWPAVALSGVPGVGVPGVGVPGAGVPGVGVPGVGVPGVGVPGVGVPGVGVPG
VGVPGVGVPGEGVPGAGSGDGMMLPWPAVALSGVPGVGVPGVGVPGAGVPGVGVPGVGVPGVGVPGVG
VPGVGVPGVGVPGVGVPGEGVPGAGVPGVGVPGVGVPGAGVPGVGVPGVGVPGVGVPGVGVPGVGVPG
VGVPGVGVPGEGVPGAGVPGVGVPGVGVPGAGVPGVGVPGVGVPGVGVPGVGVPGVGVPGVGVPGVGV
PGEGVPGAGVPGVGVPGVGVPGAGVPGVGVPGVGVPGVGVPGVGVPGVGVPGVGVPGVGVPGEGVPGA
GVPGVGVPGVGVPGAGVPGVGVPGVGVPGVGVPGVGVPGVGVPGVGVPGVGVPGEGVPGAGSGDGMM
LPWPAVALSGVPGVGVPGVGVPGAGVPGVGVPGVGVPGVGVPGVGVPGVGVPGVGVPGVGVPGEGVPGA
GSGDGMMLPWPAVALSGVPGVGVPGVGVPGAGVPGVGVPGVGVPGVGVPGVGVPGVGVPGVGVPGVGV
PGEGVPGAGSGDGMMLPWPAVALSGVPGVGVPGVGVPGAGVPGVGVPGVGVPGVGVPGVGVPGVGVPG
VGVPGVGVPGEGVPGAGSGDGMMLPWPAVALSGVPGWLDSLEFIA*

*ELP expression and purification*

Genes encoding the ELP sequences were produced using a standard iterative enzyme digestion and ligation protocol. Specifically, ELP constructs containing a single $(Q\text{-block})_1$ or $(A2V8E1)_1$ repeat in the pmA vector were obtained commercially. To generate higher-order repeats, we followed a standard recursive directional ligation strategy. Briefly, the monomeric ELP fragment was excised using PflMI and BglI and ligated (T4 DNA ligase) into a PflMI-linearized acceptor plasmid containing the same repeat to obtain the dimeric constructs $(Q\text{-block})_2$ and $(A2V8E1)_2$. This iterative excision–ligation cycle was repeated to generate the tetrameric sequences $(Q\text{-block})_4$ and $(A2V8E1)_4$. Next, the $(A2V8E1)_4$ fragment excised with PflMI/BglI was inserted into a PflMI-linearized $(Q\text{-block})_4$ vector to yield the chimeric construct $(Q\text{-block})_4$–$(A2V8E1)_4$. The resulting plasmid was again linearized with PflMI and ligated with a PflMI/BglI-excised (Q-block)4 insert to produce the final $(Q\text{-block})_4$–$(A2V8E1)_4$–$(Q\text{-block})_4$ sequence. The constructed plasmids were inserted into a pET-28a(+) expression vector and transformed into BL21 (DE3) E. coli. ELPs were expressed at 37 ˚C in TB media for 24h without the addition of inducers. The bacteria were collected by centrifugation, resuspended in 20 mM HEPES and 150 mM NaCl, and lysed by ultrasonication. The lysate was further processed by at least 3 cycles of Iterative Transition Cycling (ITC). The final ELP constructs were dialyzed in $H_2O$ for 72 hours with liquid replacement and lyophilized, aliquoted, and stored at -20 ˚C for further use.

*Large-scale ELP expression in fermenter*

ELPs were expressed at 37 ˚C for 4 hours in 10 L optimized TB media (12 g/L Tryptone, 24 g/L Yeast Extract, 0.4% v/v glycerol) with 10% TB part B, 2 mM $MgCl_2$, 10 mM L-proline and 50 μg/mL kanamycin under shaking (600 rpm) for 4 hours (no IPTG). 1mM IPTG was then added into the expression system to induce expression of the ELP for 20 hours. The cells were collected by centrifugation. The lysis and purification of ELP was the same as for the standard expression protocol (above).

*LCST measurements by UV spectrometry*

The LCST of each ELP was measured using an Evolution 260 Bio UV-Vis spectrophotometer (Thermo Scientific). 30 μM ELP and 20 mM $CaCl_2$ in HEPES were prepared in cuvettes and transferred to the spectrophotometer that had been precooled to 10 °C. Absorbance was then measured at 350 nm while the temperature was raised from 10 ˚C to 65 ˚C at a rate of 1 °C/min. The LCST was analyzed by fitting the absorbance-temperature curves using a sigmoid function.

*LC-MS analysis of ELP-peptide conjugation*

Recombinant ELPs were prepared at 5 μM and incubated with 40 μM FITC-conjugated fibrinogen γ400–411 peptide in 20 mM HEPES buffer containing 150 mM NaCl, 20 mM $CaCl_2$, and 10 μg/mL factor XIIIa at 25 °C for 1 h. Reaction mixtures were subjected to LC–MS analysis to detect and confirm formation of the covalently linked ELP–γ400–411 peptide conjugate. LC–MS was performed using a Bruker maXis 4G ESI–Q–TOF mass spectrometer coupled to a Thermo Fisher Ultimate 3000 UPLC system. Proteins were separated on a Phenomenex Jupiter C4 column (5 μm, 300 Å, 50 × 2 mm) maintained at 30 °C. The mobile phase consisted of solvent A (water with 0.1% formic acid) and solvent B (acetonitrile with 0.1% formic acid), delivered at a flow rate of 0.3 mL $min^{-1}$ using a linear gradient. Mass spectra were processed and deconvoluted using Bruker Compass DataAnalysis version 4.4.

*SDS–PAGE analysis*

SDS–PAGE was used to assess ELP purity, fibrinogen–ELP reactions, and ELP–peptide conjugation. For ELP characterization (Figure 2a), recombinant ELPs were prepared at 10 μM in 20 mM HEPES buffer, mixed with 5× SDS loading buffer containing 5% β-mercaptoethanol, and heated at 95 °C for 5 min prior to electrophoresis. For fibrinogen–ELP reactions (Figure 4a), ELPs were prepared at 15 μM and incubated with 7.5 μM fibrinogen, 1 U/mL thrombin, and 20 mM $CaCl_2$ at 37 °C for 1 h; fibrin clots were removed, and the remaining supernatants were combined with 5× SDS loading buffer and heated as above. For ELP–peptide conjugation (Figure S1), ELPs (10 μM) were reacted with 10 μM FITC-conjugated fibrinogen γ400–411 peptide in the presence of factor XIIIa (10 μg/mL) and $CaCl_2$ (20 mM) at 25 °C for 1 h before addition of SDS loading buffer and heating. All samples were resolved on 12% SDS–PAGE gels under reducing conditions. Unstained gels containing FITC-labeled samples were first imaged for fluorescence under blue epi-illumination (530/28 nm emission). Gels were then stained with Coomassie Brilliant Blue and imaged under white-light transillumination to evaluate protein loading and integrity.

*Thromboelastography (TEG) measurements*

TEG experiments of fibrinogen samples were performed using a Thrombelastograph Analyzer (Haemonetics, Boston, United States). 3.0 mg/mL of fibrinogen, 30 μM of ELP and HEPES buffer were mixed and incubated at 37 ˚C for 3 minutes, $CaCl_2$ and thrombin were then loaded to a final concentration of 20 mM and 0.2 U $mL^{-1}$ respectively. Immediately after the addition of thrombin, the solution was mixed by pipetting, and 340 μL of the sample was loaded into the TEG cup preheated at 37 ˚C. The measurement began immediately after loading the sample. The clotting onset time, α-angle, and MA values were recorded using the built in software to evaluate clotting kinetics and mechanical properties. Blood samples for TEG experiments were obtained from the Basel blood center (Blutspendezentrum, Basel, Switzerland) or extracted from ELP-treated mice by heart puncture. 300 μL of blood and the mixture of different concentrations of ELP, 20 mM of $CaCl_2$, and HEPES buffer were prewarmed in 2 separate tubes at 37 ˚C for 3 minutes. The tubes were then mixed along with 60 μL of Dade PT Innovin Recombin (Siemens Healthcare Diagnostics Inc. Newark, United States) and loaded directly into the TEG cups for measurement.

*Rheometry*

Rheometer experiments were performed using an Anton Paar MCR 302 Rheometer (Anton Paar, Graz, Austria) with a cone-plate geometry (d = 25 mm; 1˚ angle). 3.0 mg $mL^{-1}$ fibrinogen, 30 μM ELP, and HEPES buffer were mixed and incubated at 37 °C in a water bath for 3 minutes. $CaCl_2$ and thrombin were then loaded to a final concentration at 20 mM and 0.2 U/mL, respectively. Immediately after the addition of thrombin, the solution was mixed by pipetting, and 90 μL of the sample was loaded on the Peltier plate of the rheometer preheated at 37 °C. The cone plate was spun at 60 rpm for 5 seconds to ensure proper sample mixing and distribution. Silicone oil (η= 100 cSt) was then applied to the edges of the sample in order to prevent evaporation, and the clot was allowed to equilibrate for 1 h, after which time a frequency sweep was performed from 0.1 – 3 Hz (γ= 1%; previously determined to be within the Linear Viscoelastic Region (LVE) for this material).

*SEM imaging*

12 mm coverslips were cleaned in a solution of 1:1 $H_2O_2$: EtOH for 15 minutes in an ultrasonicator bath, and then washed twice with $H_2O$ twice, dried by gentle blowing under $N_2$, placed in oven at 60 ˚C for 30 min, and then stored under Ar if not used immediately. Cleaned coverslips were placed on the inverted Eppendorf tube lid inside a petri dish. SEM samples of fibrinogen were prepared by mixing 1.5 mg $mL^{-1}$ fibrinogen, 30 μM ELP and HEPES buffer and incubating at 37 °C water bath for 3 minutes. $CaCl_2$ and thrombin were then loaded to a final concentration at 20 mM and 0.1 U/mL, respectively. Immediately after the addition of thrombin, the

solution was mixed by pipetting, and 130 µL of samples were loaded on the coverslips and placed at 37 °C for 1 hour for the formation of fibrin gel. SEM samples of mouse blood were prepared by adding 150 µL of blood extract obtained by heart puncture (equal volumes of blood from male and female mice, 1:1 mixture), and 20 µL HEPES buffer containing $CaCl_2$, which led to a final mixture concentration of 20 mM. Samples were prewarmed in 2 separate tubes at 37 ˚C for 3 minutes. Samples were then mixed along with 30 µL of Dade Innovin solution and 150 µL of the mixture was loaded directly onto the coverslips. Afterwards, they were transferred to a solution of 2.5 % glutaraldehyde, and fixed at 37 ˚C for 1 h. Samples were then washed 5 times in $H_2O$ and dehydrated using increasing percentages of ethanol (25%, 50%, 75%, 100%, 100%; 10 minutes per treatment). Samples were subsequently dried by critical point drying, and then sputter-coated with platinum before being imaged in a Hitachi S-4800 SEM (Hitachi, Japan).

*Confocal fluorescence imaging*

Confocal microscopy measurements were performed using an LSM 880 ZEISS Axio Observer inverted microscope (Carl Zeiss, Germany) with a water immersion objective (C-Apochromate 40×/1.2 W korr FCS M27). For imaging experiments, fibrin clots were formed in the wells of an Ibidi µ-Slide 8 Well high Glass Bottom (Gräfelfing, Germany) from 200 µL clotting solutions consisting of 3 mg/mL fibrinogen (spiked with 1% Alexa 488 Conjugated fibrinogen, Thermo Scientific, Massachusetts, United States), 0.2 U/mL thrombin, 20 mM $CaCl_2$, and HEPES/30 µM ELP2/30 µM conELP. Fibrinogen labeled with Alexa 488 were excited using a 488 nm argon laser and fluorescence was detected in the range of 506–535 nm. ELP labelled with Alexa 594 NHS Ester (Thermo Scientific, Massachusetts, United States) were excited with a 561 nm HeNe-laser, and fluorescence was collected in the range of 603–733 nm. Each sample was scanned unidirectionally using 1024 × 1024 pixels with a Bit depth of 8 Bit. Images were processed using Fiji 2.0 software.

*Endotoxin removal from ELPs*

The Pierce™ High Capacity Endotoxin Removal Spin Columns, 1 mL (Thermo Scientific, Massachusetts, United States) was regenerated with 0.2 N NaOH and incubated overnight at room temperature, then washed with endotoxin-free 2M NaCl, water, 20 mM HEPES and 150 mM NaCl, respectively. Next, the purified ELP was loaded to the resin, and incubated with gentle end-over-end mixing at 4 °C overnight. Then the ELP was then collected by centrifugation.

*Endotoxin quantification*

Endotoxin levels were quantified using the Pierce™ Chromogenic Endotoxin Quant Kit (Thermo Scientific, Massachusetts, United States). A 96-well plate was pre-warmed at 37 °C. 50 µL of endotoxin standard dilutions, blank buffer, or samples were added to each well. The plate was kept at 37 °C, and 50 µL of the reconstituted amebocyte lysate reagent was added per well and mixed by gently tapping 10 times on the side of the plate. The plate was incubated at 37 °C for 12 min on the lysate vial. Then 100 µL of pre-warmed reconstituted chromogenic substrate solution was added to each well and incubated at 37 °C for 6 minutes. Next, 50 µL of the stop solution (25% acetic acid) was added to each well, and the optical density (OD) at 405 nm was recorded immediately.

*Animal study approval*

All animal experiments were conducted in accordance with Swiss national and Cantonal regulations on animal experimentation and animal welfare. The study was approved by the Cantonal Veterinary Office of Basel (Study #3211).

*Mouse platelets depletion by anti-CD41 antibody*

The female/male C57/B6J mice, 6-8 weeks old were treated by a single i.p. injection of an anti-CD41 antibody (Ultra-LEAF™ Purified anti-mouse CD41 Antibody, glycoprotein GPIIb, clone MWReg 30, BioLegend, San Diego, United States) at a dose of 1 mg/kg, 0.5 mg/kg, 0.1 mg/kg or 0 mg/kg diluted in 200 µL PBS. To evaluate the effect of platelet depletion by anti-CD41, whole blood samples (20 µL) were collected from the tail vein before injection and 1, 3, 6, and 24 hours after injection and diluted in 20 µL EDTA (12 mM) at a 1:1 ratio. Platelet counts were assessed using flow cytometry.

*Tail transection and ELP2 administration on platelet-depleted mice*

The female/male C57/B6J mice, 6-8 weeks old were treated with 0.5 mg/kg of anti-CD41 antibodies as described above. 24 h after antibody administration, all mice were anesthetized with isoflurane. Once anesthetized, the tip of the tail 1 mm away from the end was amputated, inflicting a controlled injury to the vessels, and initiating bleeding. Immediately after amputation, 150 mg/kg, 50 mg/kg, or 0 mg/kg of ELP solution were i.p. injected. Immediately after injection, the mice were placed on a heated mat and their tails were immersed in 15 mL tubes with saline (37°C, 0.9% NaCl). Blood was collected over two time periods of 30 minutes each. After each 30 minute period, the collection tube was removed and placed on ice. Total blood loss amount was determined by hemoglobin measurement in collected samples using the Soret absorbance band at 405 nm. A standard curve derived from defined blood volumes was used to calculate the volume of lost blood in µL. After 60 minutes, the mice were euthanized via $CO_2$ asphyxiation and blood was collected by cardiac puncture for further analysis.

*Quantification of ELP in blood post administration*

Cyanin 7.5 NHS-Ester (Lumiprobe Corporation, Maryland, USA) labelled ELP2 was intraperitoneally injected at a concentration of 150 mg/kg. 10 µL of blood was then taken through a tiny cut in tail 5 min/30 min/1 h/3 h/6 h/24 h post administration. 780 nm of excitation light and 808 nm of emission light was adopted to measure the fluorescent intensity of cyanin 7.5 labelled on ELP. A standard concentration of cyanin 7.5-ELP, ranging from 7.5 g/mL to 7.5 mg/mL in blood, was used to calibrate the signals measured in the plate reader.

*In Vivo Imaging*

Mice were anesthetized using isoflurane and imaged by an in vivo Imager (IVIS Lumina Series III, PerkinElmer). Data was acquired and analyzed using the Living Image Software.

*ELISA quantification of serum IgG/IgM level in mice after ELP administration*

Mouse IgM Uncoated ELISA Kit (Invitrogen, #88-50470) and Mouse IgG (Total) Uncoated ELISA Kit (Invitrogen, #88-50400) were used to detect antibody IgM/IgG level in mice. Corning Costar 9018 ELISA plate was coated with 100 µL/well of capture antibody in coating buffer overnight at 4 °C. The plate was then blocked with blocking buffer at room temperature for 2 hours. 100 µL of reconstituted IgG/IgM standard or prediluted serum samples were added to each well and incubated at room temperature for 2 hours. Substrate solution and stop solution was added sequentially to each well. Absorbance signal was read at 450 nm.

### Supporting Figures

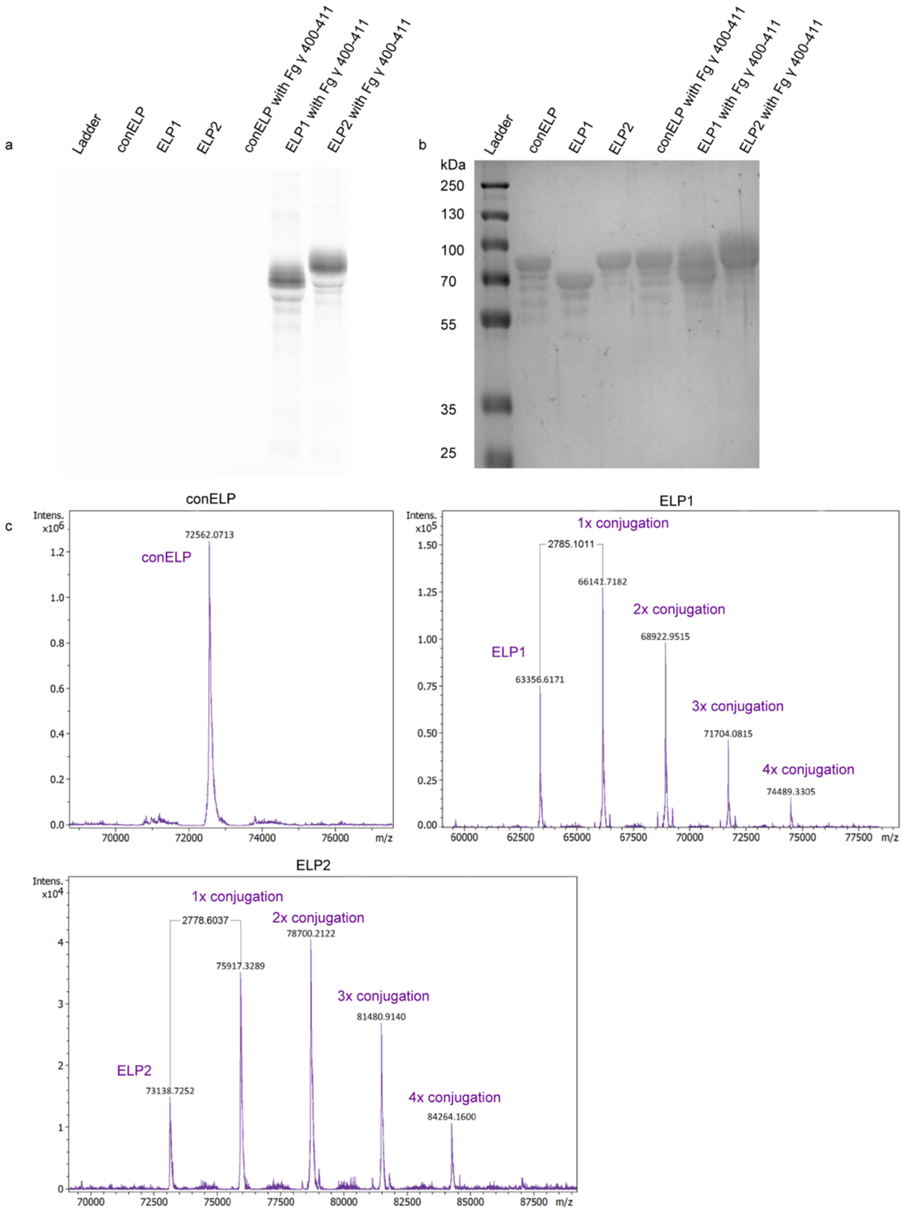


**Figure S1.** SDS–PAGE analysis of Q-block-ELP and fibrinogen γ 400-411 peptide crosslinking reactions. Q-block-ELPs were incubated with FITC-conjugated fibrinogen γ 400-411 peptide in the presence of factor XIIIa and $CaCl_2$ for 1 h at 25 °C. (a) Following electrophoresis, the gel was first imaged for FITC fluorescence under blue epi illumination with a 530/28 nm emission filter. (b) Next, the gel was stained with Coomassie Brilliant Blue and imaged under white light transillumination. (c) Reaction products were analyzed by LC-MS, which confirmed formation of the covalently linked Q-block-ELP with the fibrinogen γ400–411 peptide. ELP1 and ELP2 are theoretically able to be conjugated up to eight times. We can observe most conjugations occurring once or twice, respectively. The maximal observable degree of modification for both ELPs was four.

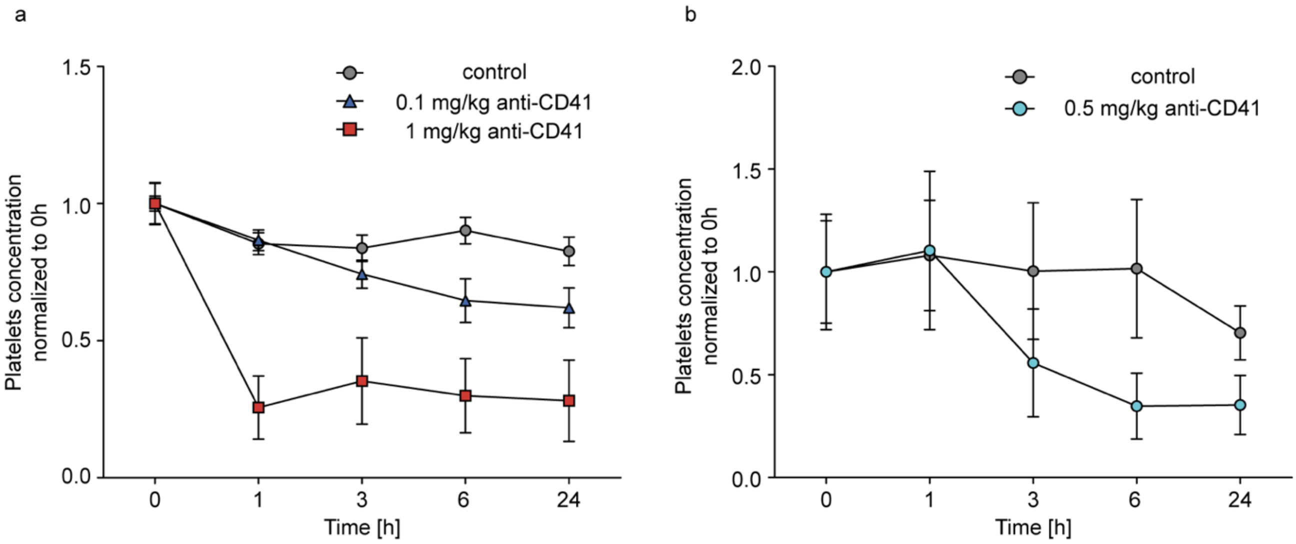


**Figure S2**. Mouse platelet depletion by anti-CD41. The concentration of platelets was normalized to its original level before anti-CD41 injection (0h). (a) Anti-CD41 concentrations: red, 1 mg/kg; blue, 0.1 mg/kg; grey, 0 mg/kg. (b) Anti-CD41 concentrations: blue, 0.5 mg/kg; grey, 0 mg/kg.

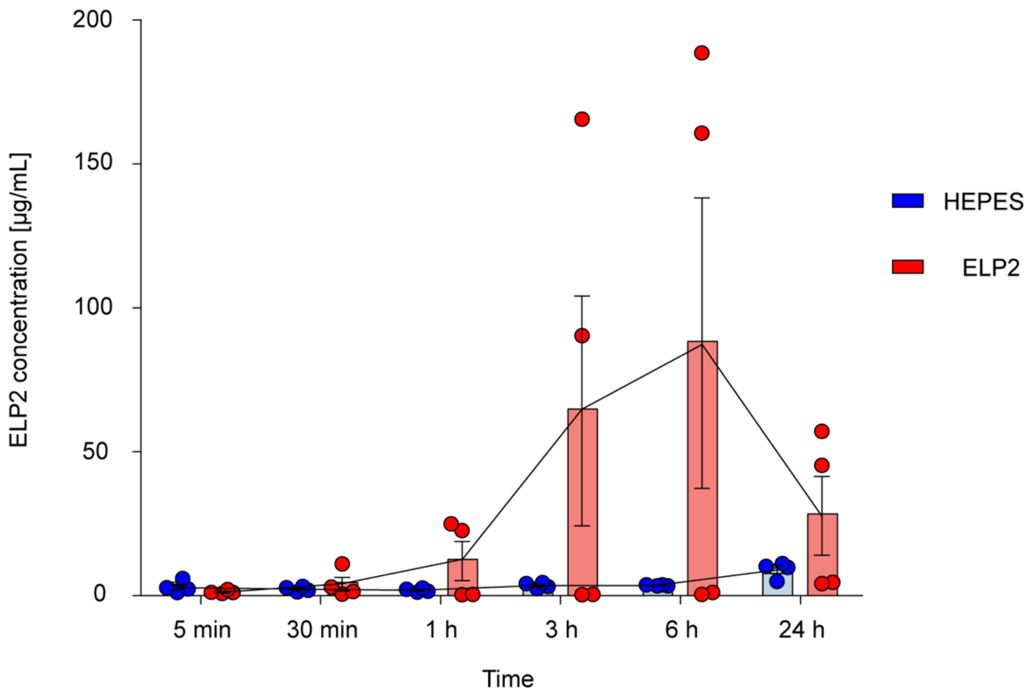


**Figure S3**. ELP2 concentration in blood after administration. Concentration of ELP2 in mouse blood was measured 5 min/30 min/1 h/3 h/6 h/24 h post administration. 4 mice were injected with 150 mg/kg of Cyanin7.5-labelled ELP (red). 4 mice were injected with HEPES as negative control (blue).

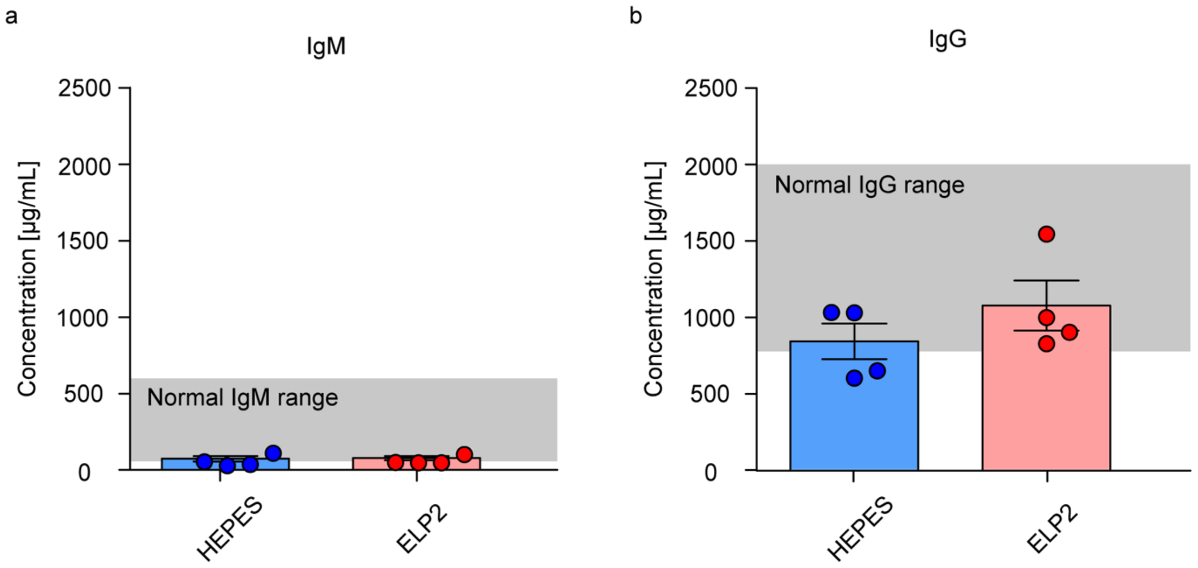


**Figure S4**. In vivo antibody levels in mice after subcutaneous injection of ELP2. Serum concentrations of total IgM (a) and IgG (b) 7 days post-injection with HEPES (blue)/ELP (red). The grey band indicates the reported physiological ranges of total serum IgM (0.1-0.6 mg/mL) and IgG (0.8–2.0 mg/mL) in adult C57BL/6 mice.

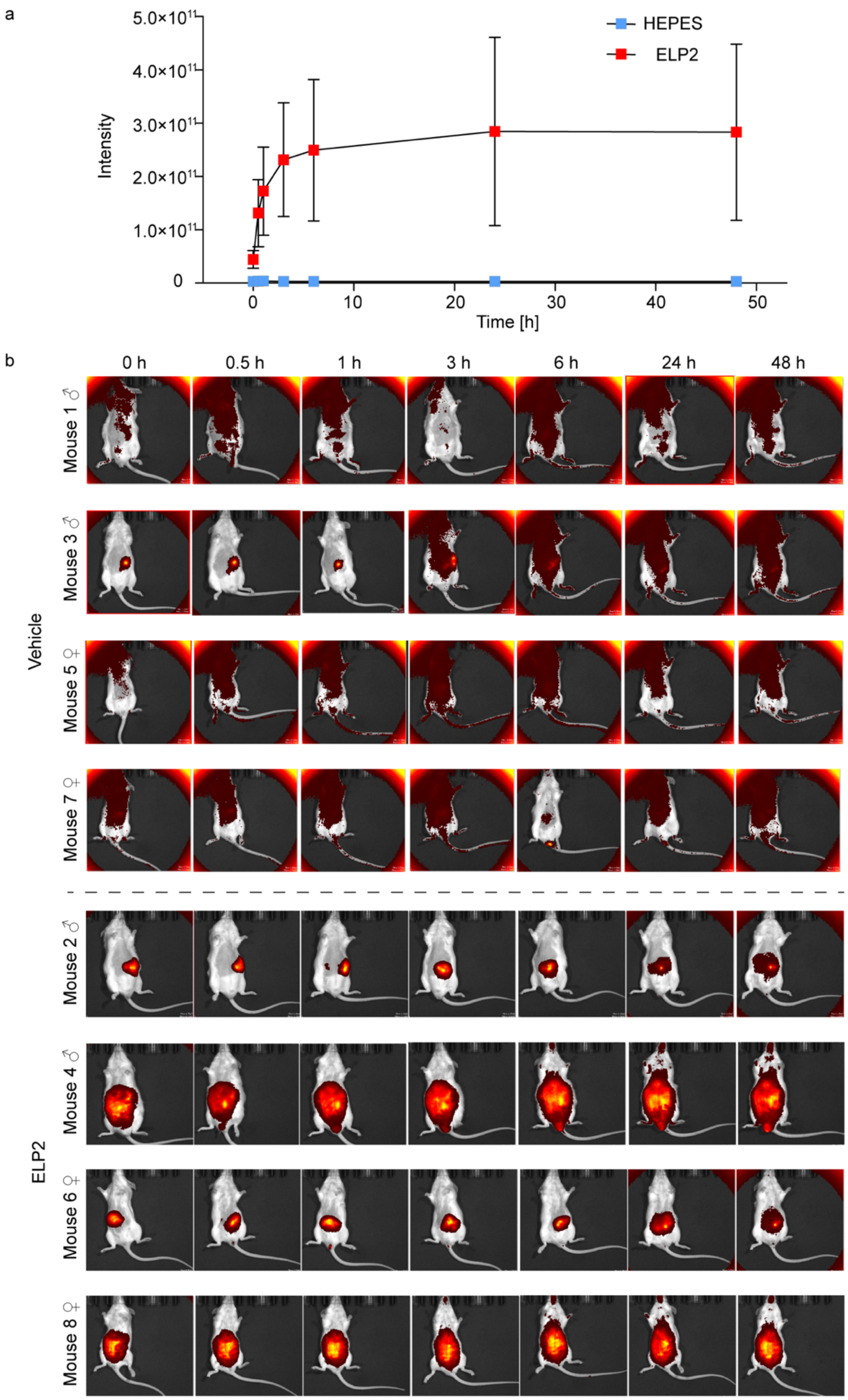


**Figure S5.** In vivo biodistribution of ELP2. (a) Quantification of the fluorescence intensity, with the error bars showing mean ± s.e.m. (n = 4 at each time point). (b) In vivo fluorescence images of the wild-type mice treated with cyanin 7.5-labelled ELP2 at different time intervals.

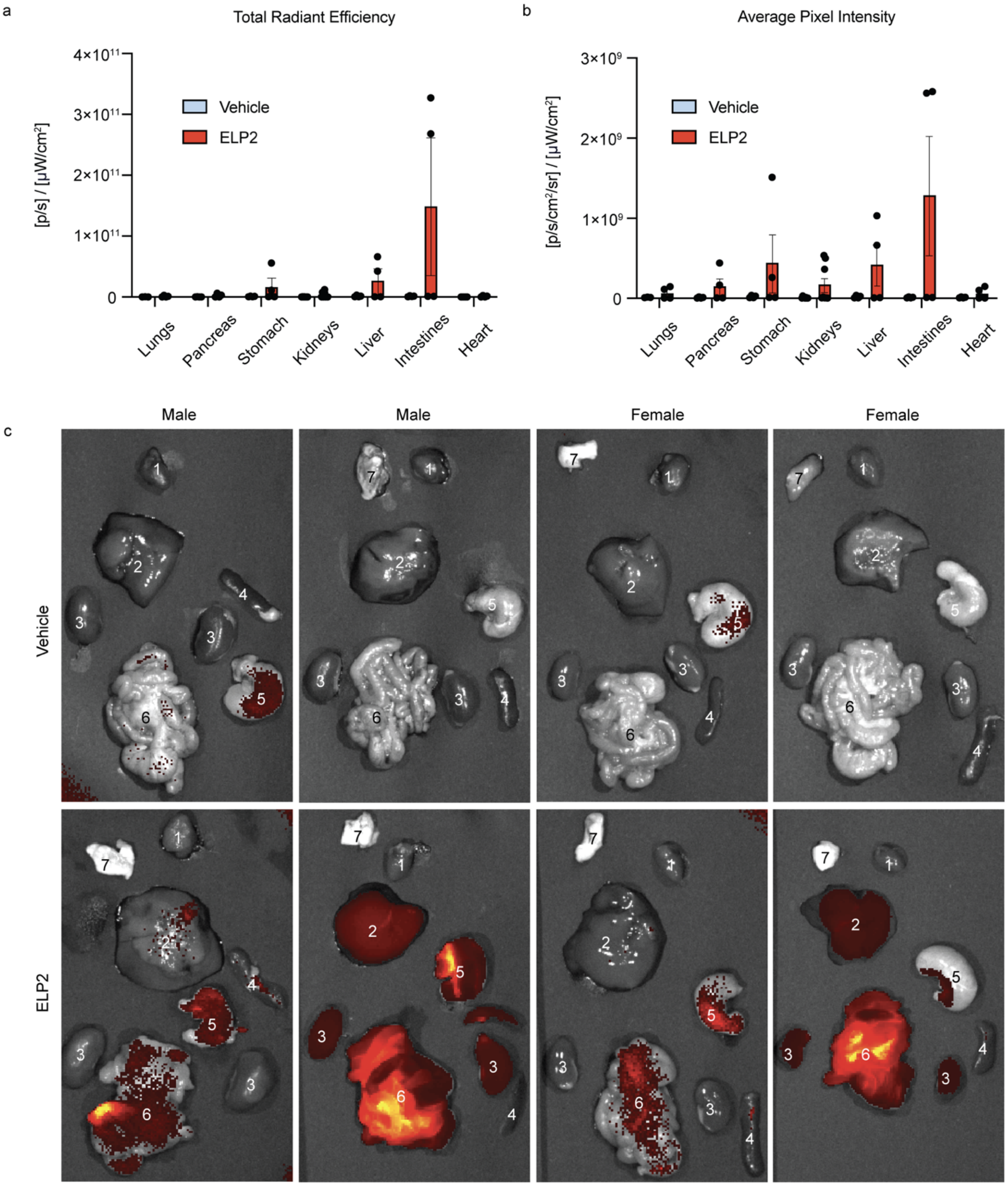


**Figure S6.** Biodistribution analysis of ELP2 in organs. Quantification of the (a) total radiant efficiency and (b) average pixel fluorescence intensity in different organs, with the error bars showing mean ± s.e.m. (n = 8 for kidneys, and n = 4 for other organs) 48 hours post i.p. injection of cyanin 7.5-labelled ELPs. (c) Fluorescence images of organs from wild-type mice treated with cyanin 7.5-labelled ELPs. The organs are labeled with numbers 1-7 for identification (1: heart, 2: liver, 3: kidneys, 4: pancreas, 5: stomach, 6: intestines, 7: lungs).

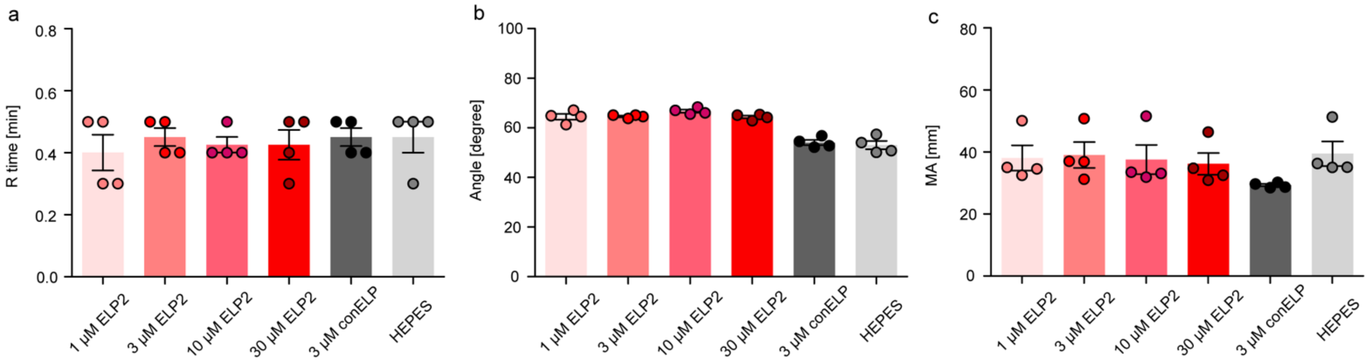


**Figure S7.** The effects of different concentrations of ELP2/conELP/HEPES on the kinetics and mechanical stability of human blood clots with normal platelet counts, measured by thromboelastography. (a) Clotting onset time (R time). (b) α-angle. (c) maximum amplitude (MA).

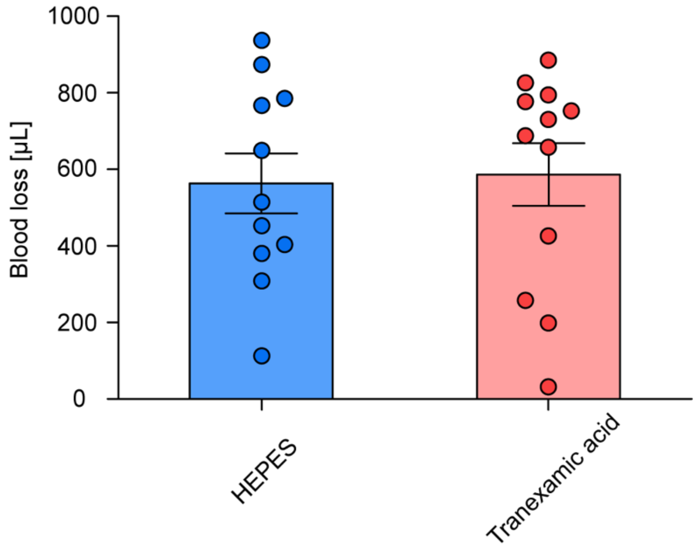


**Figure S8.** Tail bleeding assays performed on platelet-depleted thrombocytopenic mice showed no change in blood loss between animals treated with tranexamic acid vs. HEPES buffer (vehicle).
